# High regional variation of spotted lanternfly suitability in New York wine-growing regions signals uneven vineyard risk

**DOI:** 10.1101/2025.05.28.656630

**Authors:** Jaclyn A. Eller, Kaitlin M. Gold, Sara E. Emery

## Abstract

The spread of the spotted lanternfly (*Lycorma delicatula*) is facilitated by its ecological flexibility and generalist diet breadth. Among 100 other plants, spotted lanternfly feed on grapevine, so high vineyard density could make suitable habitat even more attractive. This invasive polyphagous insect threatens New York viticulture, a >$6 billion industry, but the local risk remains uncertain. New York state may include boundary conditions defining the edges of environmentally suitable habitat.
Using 16,028 spotted lanternfly observations, a species distribution model was developed for the northeastern U.S. using climatic, geographic and human influence variables. Suitability was modeled at 1km under both current (2011-2040, RCP 7.0) and worsening near-future (2041-2070, RCP 8.5) climate scenarios. Suitability models were also developed for four common tree hosts (tree -of-heaven, black walnut, red maple and sugar maple), to evaluate if combined models that include established plant host suitability improve estimates of spotted lanternfly occurrence. Kernel Density estimates of vineyard density were used to add a maximum of 10% risk.
Combining the spotted lanternfly suitability with tree-of-heaven suitability improves estimates of spotted lanternfly occurrence. Human influence and humidity were important for defining suitability. Frost change frequency increased in importance under near-future climate change conditions. On Long Island, 83% vineyards are at very high risk for spotted lanternfly (>75% suitability). Across New York state, 33% of all vineyards are classified as very high risk. Under future climate change conditions, the number of vineyards at very high risk of spotted lanternfly increases by 7% in the Finger Lakes but decreases by 22% on Long Island.
This model improves estimates of spotted lanternfly potential distribution by integrating suitability of its established host tree-of-heaven. Estimating potential expansion under near-future climate conditions provides information on variables that define the environmental limits of suitability. Quantifying the local risk of vineyards to spotted lanternfly at the 1km scale creates an important tool for allocating resources to mitigate and manage this pest.

**Open research statement:** Most data are already published and publicly available, with those items properly cited in this submission (GBIF species observations, CHELSA CMIP6 environmental variables, USGS topography, NASA remote sensing). However, observations of spotted lanternfly from the New York State Department of Agriculture and Markets (N=10,144) and specific vineyard locations from the New York Wine & Grape Foundation (N=6,651) are considered sensitive information and are not shared here. Researchers interested in these datasets may reach out to these organizations to request access.

## Introduction

Invasive species disrupt natural and agricultural ecosystems to cause cascading environmental harm, including shifts in community biodiversity and economic losses (Xu et al. 2006; Oliveira et al. 2013; Boivin et al. 2016). Climate change has exacerbated invasive species spread worldwide (Finch et al. 2022; Worm et al. 2024). The spotted lanternfly (*Lycorma delicatula* (White) [Hemiptera:Fulgoridae]), a univoltine polyphagous planthopper native to China, was first detected in the United States (U.S.) in Pennsylvania in 2014 (Dara et al. 2015, Du et al. 2021, Uyi et al. 2021). Adults lay eggs on materials that are often moved through human activities, increasing the rate of population spread (Elsensohn et al. 2024; Strömbom et al. 2024; Ladin et al. 2023). Although the spotted lanternfly is not considered a long-distance flyer, it readily hitchhikes by laying egg masses on various outdoor surfaces such as lumber, stone, and vehicles, facilitating accidental transport to novel regions (Cooperband et al. 2019; Myrick and Baker 2019; Ladin et al. 2023a). Spotted lanternfly also persists under a wide range of climatic conditions (Keena et al. 2023). High phenotypic plasticity may be as important as genetic diversity in maintaining fitness in novel habitats and under changing climatic conditions (Ghalambor et al. 2007). Species with extensive geographic distributions often demonstrate high phenotypic plasticity (Bennett et al. 2019; Valladares et al. 2014), improving success in novel regions of recent invasion and under novel conditions due to climate change (Thompson et al. 2021).

The spotted lanternfly can survive and develop on a single plant host species, but feeding on multiple host species enhances fitness and egg production (Laveaga et al. 2023). The spotted lanternfly is highly polyphagous, feeding on over 100 plant species across 33 families (Barringer and Ciafré, 2020). The spotted lanternfly can be found in both forests and cultivated crops across different life stages (Booth et al. 2025; H. Leach and Leach 2020; A. Leach and Leach 2020). Spotted lanternfly communication and attraction is linked to a combination of host plant odors (e.g., methyl salicylate), species-specific pheromone blends, and substrate vibrations (Cooperband et al. 2019; Derstine et al. 2020; Cooperband and Murman 2024). Adults demonstrate preference for tree-of-heaven (*Ailanthus altissima* (Mill.) Swingle), a widespread invasive deciduous tree introduced to the U.S. in 1784 (Soler and Izquierdo 2024). When tree of heaven is limited or unavailable, alternate hosts, such as red maple and sugar maple (*Acer* spp.), black walnut (*Juglans* spp.), and grapevines (*Vitis* spp.), can support all life stages (Cooperband et al. 2019; Murman et al. 2020; Laveaga et al. 2023; Booth et al. 2025). Spotted lanternfly presents a high economic risk to many crops, especially cultivated grapes (*Vitis labrusca*, *Vitis vinifera* and *Vitis vinifera* interspecific hybrids) (A. Leach and Leach 2020; Murman et al. 2020; Huron et al. 2022). Spotted lanternfly exposed to a mixed diet of grapevine and tree-of-heaven resulted in greater nymphal development, egg production, and body mass compared to those fed on either species alone (Laveaga et al. 2023, Nixon et al. 2022).

Excessive spotted lanternfly feeding reduces grapevine plant health and reduces winter survival (Harner et al. 2022; Lavely et al. 2022). Spotted lanternfly-invaded vineyards have reported yield loses >90% and vine death after heavy infestations (H. Leach and Leach 2020; Murman et al. 2020; Urban 2020). New York state is the third largest producer of wine and juice grapes in the country and generates $6.65 billion in economic activity (Dunham & Associates 2022). Vineyards are primarily located in four regions across the state: Lake Erie, the Finger Lakes, the Hudson Valley, and on Long Island. Although, the exact economic impact of spotted lanternfly on New York’s grape industry is unknown, a recent study has estimated that losses due to spotted lanternfly could be as high as $1.5 million in the first year after infestation (Pinto et al. 2025). Since its first confirmed report in New York state in 2017, researchers and stakeholders have shown increasing interest in quantifying spotted lanternfly responses to spatiotemporal climatic variability (Lewkiewicz et al. 2024; Barker et al. 2025).

Modeling strategies used for identifying spotted lanternfly habitat suitability include predictive and process-based frameworks and correlative climatic suitability models at regional and global scales (Wakie et al. 2019; Cook et al. 2021; Jones et al. 2022; Strömbom et al. 2024; Zhao et al. 2024; Barker et al. 2025). Information is rarely available at a fine enough resolution to allocate scarce resources towards targeted mitigation and management efforts. Additionally, most models use averages of historical climatic conditions (e.g., CMIP3 and CMIP5), resulting in potentially inaccurate establishment risk because of climate change. Many species distribution models do not incorporate the broad diversity of variables, beyond climatic ones, that we know to be important in defining a species niche (Mod et al. 2016). Remote sensing imagery reflects real-time data on land use, vegetation health, and climatic conditions and can improve the accuracy of ecological models (Zhang et al. 2017). High-resolution remote sensing techniques can identify ecological conditions that support pest populations, allowing vulnerable areas to be identified (Wang and Hu 2021; Emery et al. 2024). Combining fine-resolution remote sensing data with near-future climate change projections is especially relevant for spotted lanternfly because of its broad diet breadth and because identifying environmental tolerance limits can clarify high risk areas, increasing the likelihood of early detection (Koch 2021; Hao et al. 2024).

In anticipation of the risk to the New York grape growing industry, the New York State Department of Agriculture and Markets (NYSDAM) initiated a statewide spotted lanternfly monitoring effort in 2020 to track the invasion progress from the southeastern part of New York state that has since collected over 10,000 observations of spotted lanternfly. Further citizen science efforts from the Global Biodiversity Information Facility (GBIF) in previously invaded neighboring states offered 5,884 additional observations (GBIF.org 2026). Much of the Species Distribution Model literature uses MaxEnt (Bradie and Leung 2017; Fourcade et al. 2014; Phillips et al. 2006), including Species Distribution Models developed specifically for spotted lanternfly (Huron et al. 2022; Namgung et al. 2020; Wakie et al. 2019). However, problems have been identified with Species Distribution Models built with MaxEnt, and those built with random forest were shown to perform better overall (Zhao et al. 2022). The phenotypic plasticity of spotted lanternfly, the widespread distribution of established plant hosts and the potential for this invasive pest to cause economic damage, all make this an ideal system to develop a fine-scale landscape-informed correlative species distribution model using Random Forest.

The goal of this study was to quantify the risk of spotted lanternfly to New York viticulture now and in the near-future by combining vineyard density with spotted lanternfly. Since spotted lanternfly is currently invading New York state, a species distribution model developed from occurrence records may not capture the full range of environmental tolerances of this species. Using the habitat suitability of established plant hosts across the range of spotted lanternfly occurrence records could offer additional predictive power. Many trees that are host plants to spotted lanternfly are high in Methyl salicylate and this compound has been shown to be attractive to spotted lanternfly (Cooperband et al. 2019; Booth et al. 2025; Laveaga et al. 2023; Murman et al. 2020). We integrate tree-host habitat suitability maps with spotted lanternfly habitat suitability to evaluate whether combining tree host suitability maps with the spotted lanternfly suitability map improves estimates of spotted lanternfly presence across the northeastern U.S. Approximately 8,000 observations of each of the four alternate tree-host species, tree-of-heaven, black walnut, red maple, and sugar maple, were available and downloaded from GBIF to develop alternate plant host suitability maps for New York and other states in the northeastern U.S. where spotted lanternfly has been observed (GBIF.org 2026a). We use weather variables representing current climatic conditions (2011-2040; RCP 7.0), and under near-future worsening emissions (2041-2070; RCP 8.5) scenarios to estimate spotted lanternfly and alternate tree hosts suitability. Vineyard density was included as an additive risk factor, since grapevines are another preferred host plant and in some grape-growing regions vineyards are highly clustered.

Specifically, we sought to:

1. Create a spotted lanternfly habitat suitability for the northeastern US with a correlative species distribution model, using large observational datasets, climatic, geographic and remote sensing variables under current and near-future climatic conditions.
2. Assess if integrating long-established alternate tree-host species distribution maps increases accuracy of predicting spotted lanternfly occurrence.
3. Identify variables that define spotted lanternfly habitat suitability under current climatic conditions compared to those in the near-future under a worsening climate change scenario.
4. Quantify the potential risk of spotted lanternfly for New York vineyards at a 1km scale using vineyard density as an additive risk factor.

## Materials and Methods

### Data sources

The New York Wine and Grape Foundation shared 6,651 vineyard locations across the state (**Fig. 1**). Georeferenced observations of the spotted lanternfly in New York state were obtained from the NYSDAM (N=10,144). Observations of spotted lanternfly in other northeastern states (Pennsylvania, Virginia, West Virginia, Ohio, Maryland, New Jersey, Connecticut, Massachusetts, New Hampshire, Rhode Island, Vermont, and Maine) were taken from GBIF (N= 5,884), resulting in 16,028 observations across all life-stages (**Fig S1**) (GBIF.org 2026a). Observations represent occurrence records across all life stages. Validated observations of spotted lanternfly from the NYSDAM were available only for New York state. Since New York state was the focus of the study and is an area actively being invaded, these higher observations were prioritized for inclusion in the dataset over GBIF observations available for New York state to avoid double-counting.

**Fig. 1:**
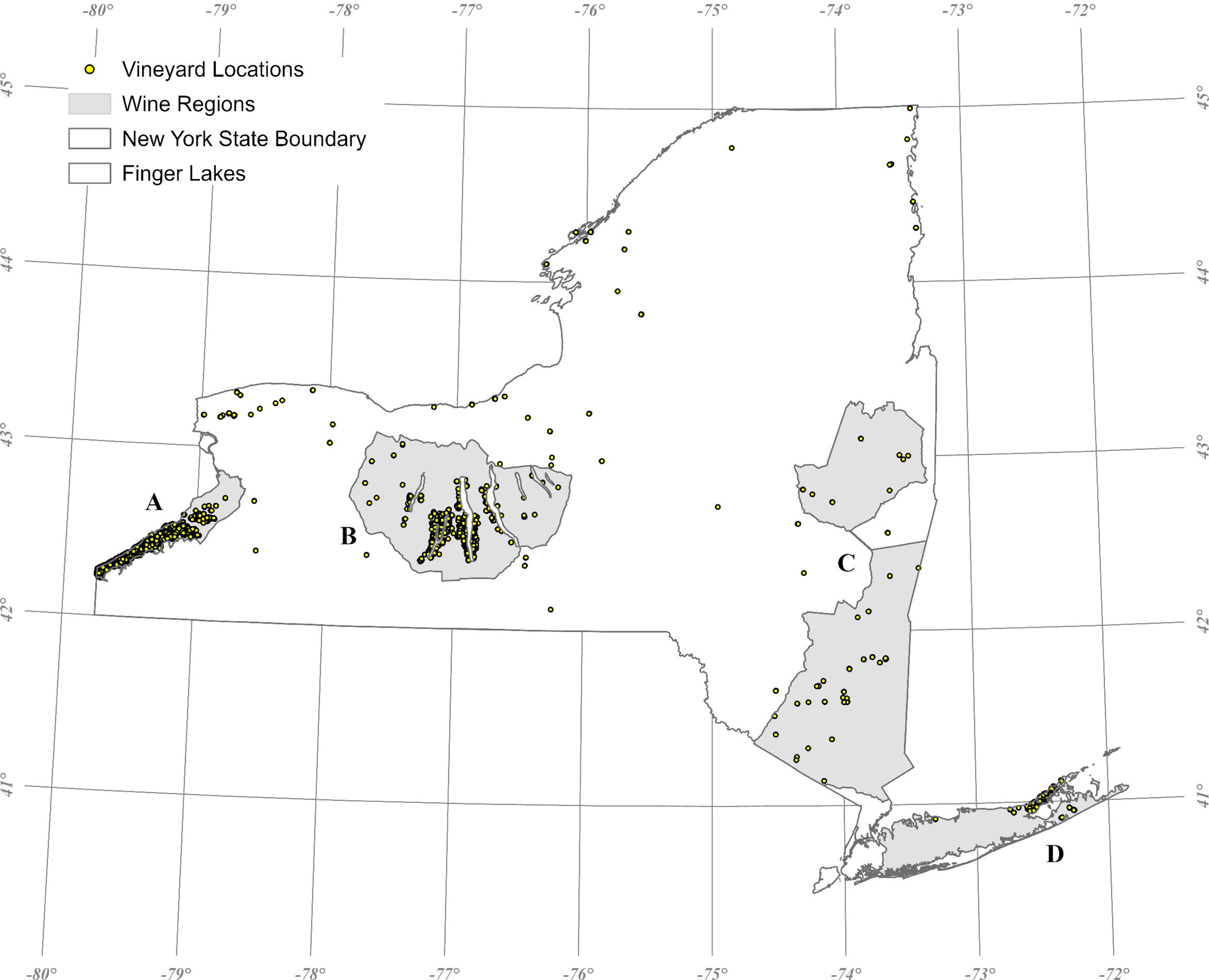
Map of 6,651 vineyard locations across New York state with its wine regions highlighted in gray: A) Lake Erie (N=3,700), B) Finger Lakes (N=2,280), C) Hudson Valley (N=118), and D) Long Island (N=381), where *N* indicates the number of vineyards within each region.

Georeferenced observations of five host-tree species known to be preferred by spotted lanternfly, *Ailanthus altissima* (Mill.) Swingle (tree of heaven), *Juglans nigra* L. (black walnut), *Acer rubrum* L. (red maple), *Acer saccharum* Marshall (sugar maple), were obtained from GBIF (GBIF.org 2026). Occurrences of tree of heaven (N = 7,756), black walnut (N = 8,334), red maple (N=8,370), and sugar maple (N=8,517) in New York as well as across northeastern states (Pennsylvania, Virginia, West Virginia, Ohio, Maryland, New Jersey, Connecticut, Massachusetts, New Hampshire, Rhode Island, Vermont, and Maine), were obtained from GBIF (GBIF.org 2026b,c,d,e). All GBIF observations were extracted and cleaned using R package *CoordinateCleaner* (Zizka et al. 2019) in R version 4.5.1 (R Core Team 2025) (Supplementary Methods 1.1).

### Climatic Variables

To model habitat suitability for spotted lanternfly and the four tree species, daily and monthly mean climatic variables were downloaded from CHELSA CMIP6 ISIMIP3 version 2.1 (Karger et al. 2017). Focusing primarily on GFDL-ESM4 datasets, isothermality, frost change frequency, temperature seasonality, precipitation seasonality, and mean monthly precipitation of the coldest quarter were assessed under both current (2011-2040; SSP 370, RCP 7.0) and near-future (2041-2070; SSP 585, RCP 8.5) conditions (**Table S1**). These variables are from Phase 6 of the Coupled Model Intercomparison Project (CMIP6). Variables with a correlation above 0.8 were not included in any Species Distribution Model.

Humidity is a critical environmental variable that is frequently omitted from species distribution models, despite its physiological relevance for ectotherms (Brown et al. 2023). Monthly near-surface relative humidity from the National Center for Atmospheric Research (NCAR) Community Climate System Model 4 run Phase 5 of the Coupled Model Intercomparison Project (CCSM4 CMIP5) was included as an additional climatic variable. Monthly humidity was unavailable for CMIP6. To ensure consistency between CMIP5 and CMIP6, R version 4.5.1 (R Core Team 2025) was used to extract the near-surface humidity estimates for years corresponding to the current (2011-2040, RCP 7.0) and near-future (2041-2070, RCP 8.5) climatic scenarios. Due to its potential importance, near-surface relative humidity was downscaled from 1 km to 30 m, following the framework implemented by Alves *et al*. (2025) (Supplementary Methods 2.1).

### Geographic Variables

All geographic variables used in habitat suitability models are detailed in **Table S1**. Mean 30-arc sec elevation and topographic position (**Table S3**) covariates were obtained via U.S. Geological Survey (GMTED 2010). The soil texture was obtained from the U.S. Department of Agriculture (Hengl 2018) (**Table S4**).

Vegetation phenology can be utilized to detect and map the spatial distribution of plant species in remote sensing applications (He et al. 2011; Rocchini et al. 2015). The Normalized Difference Vegetation Index (NDVI), which characterizes crop vigor and biomass during key phenological stages, has been widely used to measure vegetation productivity and improve overall mapping precision (Farooque et al., 2023; Leitão & Santos, 2019; Wang et al., 2025) (**Table S5**). The Harmonized Landsat Sentinel-2 was used to calculate the NDVI, at 30m spatial resolution, to characterize New York’ s autumn season between August 1, 2024, and November 1, 2024, (**Supplementary Methods 3.1**).

Tree canopy cover can be a useful proxy for quantifying canopy density and spatial configuration that influence host tree occurrence, edge structure, and microclimatic conditions relevant to insect establishment and movement (A. Leach and Leach 2020; Madalinska et al. 2022; Booth et al. 2025; Martinuzzi 2009). The Landsat Tree Canopy Cover dataset (Sexton et al. 2013), at 30 m spatial resolution, was also included to predict spotted lanternfly habitat suitability (Gorelick et al., 2017).

To account for the importance of human-assisted transportation corridors in the spread of spotted lanternfly (Elsensohn et al. 2024), we included the human influence index dataset (**Table S6**), at 1km resolution, was obtained from the NASA Socioeconomic Data and Applications Center (Venter et al. 2024) (**Supplementary Methods 3.2**).

### Spotted lanternfly habitat suitability

Random Forest classification was used to identify important variables defining habitat suitability for spotted lanternfly and alternative tree hosts under current (2011-2040; SSP 370, RCP 7.0) and near-future worsening (2041-2070; SSP 585, RCP 8.5) climate conditions (Chen et al. 2021). All models were developed at the 1km scale estimating suitability for the period 2011-2040 and using climate variables drawn from SSP 370, RCP 7.0, considered reflective of current conditions, or for the period 2041-2070 using climate variables drawn from SSP 585, RCP 8.5, representing near-future climate conditions. R packages SDMtune (Vignali et al. 2020), flexsdm (Velazco et al. 2022), and blockCV (Valavi et al. 2019), *randomForest* (Breiman 2001), and *pdp* (Greenwell 2017) were implemented in R 4.5.1 (R Core Team 2025) with default settings unless stated otherwise. All habitat suitability maps were projected to EPSG:4326 and illustrated in ArcGIS Pro (v3.3) (Esri 2024).

Occurrence datasets for spotted lanternfly (**Fig S1**), tree of heaven, black walnut, red maple, and sugar maple represented only presence data. True absence data were unavailable, therefore pseudo-absences were generated by randomly sampling an equal number of species observations for each focal taxon (Supplementary Methods 4.1). This resulted in a balanced presence-absence dataset for training and validation with spotted lanternfly= 31,996, tree of heaven= 15,512, black walnut= 16,668, red maple= 16,740, and sugar maple= 17,034 observations and pseudo-observations. A K-fold cross-validation approach with spatial blocks was implemented to minimize spatial autocorrelation and ensure an independent dataset for validation (Soley-Guardia et al. 2024) (Supplementary Methods 4.2). This approach is recommended for spatially structured data, regardless of the degree of clustering, spatial autocorrelation, or species abundance class (Mushagalusa et al. 2024).

Partial dependence plots were calculated to quantify the marginal relationship between individual predictor variables and the predicted probability of species presence derived from the Random Forest model (Cutler et al. 2007; Nayak et al. 2022). Variable importance was assessed using the Mean Decrease Gini Index, defined as the sum of all reductions in Gini impurity attributable to a given variable across the Random Forest splits, normalized by the number of trees (Calle and Urrea 2011; Grabska et al. 2019). Model performance was evaluated by generating a confusion matrix, including accuracy, sensitivity, specificity, positive prediction value, and negative prediction value. Final metrics for alternate tree-host species can be found in the Supplementary Materials 4.3-4.6.

To assess whether integrating distribution maps of previously established tree-hosts improves model fit of spotted lanternfly suitability, model fit of suitability output maps were compared. The fit of the spotted lanternfly suitability map without tree-host integration was compared to a spotted lanternfly suitability map that was integrated with a suitability map 1) tree-of-heaven, 2) black walnut, 3) red maple, 4) sugar maple, and 5) all tree-hosts combined. Data integration of combined maps was approached by selecting either the mean or the maximum suitability value, representing a worst-case-scenario, for each pixel in the map (G. Zhang et al. 2020; Boulesnane Guengant et al. 2025). Suitability output of each model was evaluated against spotted lanternfly presence-only occurrences. Pseudo-absences were excluded from this analysis due to potential bias in georeferenced data (Chapman et al. 2019). For each presence-only occurrence point, the predicted pixelwise suitability value was extracted to assess whether observed presences corresponded to areas with high modeled spotted lanternfly suitability. Model performance was evaluated using the Receiver Operating Characteristic Area Under the Curve (ROC AUC), which quantifies how well the model estimates occupied sites, with values closer to 1 indicating better suitability classification Zhang *et al*. (2020). Precision recall area under the curve (PR AUC), along with their corresponding standard deviations are also reported (**Table S9; S10; S11; S12**).

### Vineyard risk

A vineyard density spotted lanternfly risk multiplier map was developed from the 6,651 vineyard locations in New York using a hybrid adaptive bandwidth Kernel Density Estimate. A hybrid method combining an adaptive bandwidth by varying the kernel size and density-weighting by varying kernel density is considered superior (Carlos et al. 2010). Resolution was set at 1km to match the scale of species distribution models developed in this analysis, bandwidth was set at 20km and a density multiplier of 2.0 was implemented. The pairwise Euclidean distance to the nearest 10 neighbors and the mean distance was used to calculate local vineyard density. Values were normalized between 0-0.10 (0-10%). Ten percent was selected as the additive risk factor since this is the approximate increase in survival when spotted lanternfly is fed on a mixed diet of grape vines and tree-of-heaven compared to a diet of tree-of-heaven alone (Laveaga et al 2023). The values calculated from this adaptive Kernel Density Estimate of vineyard density was added to the combined habitat suitability map under both current (2011–2040; RCP 7.0) and near-future (2041–2070; RCP 8.5) climate conditions. Locations with the highest vineyard density added a maximum of 10% risk to the combined spotted lanternfly tree-of-heaven suitability values.

## Results

### Spotted lanternfly habitat suitability

The spotted lanternfly model under current (2011-2040; RCP 7.0) climatic conditions demonstrated high accuracy (92.3%), positive prediction value (92.0%), and negative prediction value (92.6%) (**Table S7**). The spotted lanternfly model performed well with an ROC AUC value of 97.8% under current conditions (**Table S8**). The Hudson Valley and Long Island regions appear to have the highest suitability, though under near-future climate change conditions the Hudson Valley appears to increase in suitability while Long Island declines (**Fig. 2, Fig S2**).

**Fig. 2:**
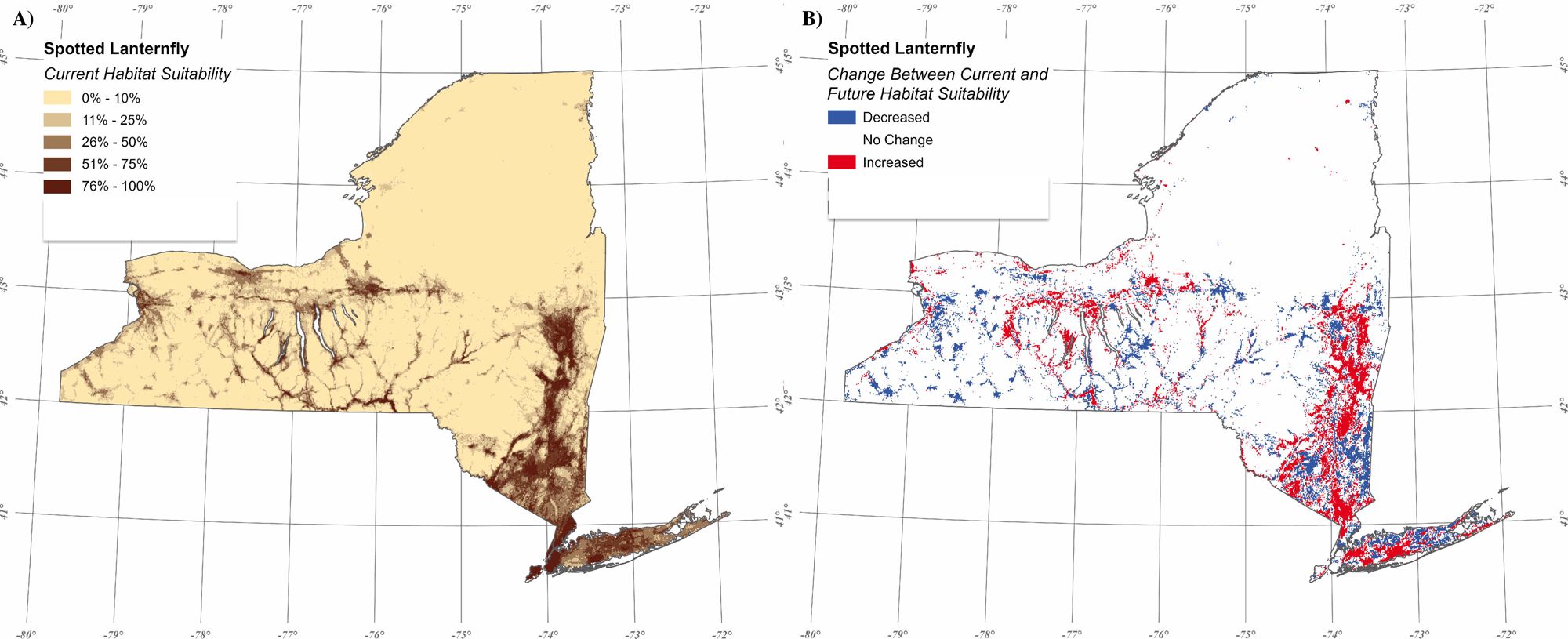
Habitat suitability for spotted lanternfly across New York state is shown in A) under current conditions (2011-2040; RCP 7.0), and in B) the change between current (2011-2040; RCP 7.0) and near-future (2041-2070; RCP 8.5) habitat suitability for spotted lanternfly across New York state is shown.

The most influential explanatory variables for modeling the spotted lanternfly between both current and near-future climatic conditions were human influence index, near-surface relative humidity, temperature seasonality and isothermality (**Fig S3**). Isothermality is an indicator of temperature fluctuation over a 24-hour period relative to annual temperature fluctuation. Under current conditions, isothermality between 20% and 25% of the difference between the hottest and coldest months are associated with higher spotted lanternfly suitability (**Table S9, Fig S4**). The highest suitability for spotted lanternfly habitats under current and future conditions is characterized by areas with low to moderate human influence and near-surface relative humidity between 72% and 80% and tree canopy cover <89% (**Table S9, Fig S4**). Spotted lanternfly suitability is positively correlated with higher frost change frequency, expected to increase in the near-future (**Table S1**). In the near-future, under worsening climate change conditions, frost change frequency becomes a more important variable defining habitat suitability for spotted lanternfly. Near surface relative humidity and tree canopy cover also become more important. The importance of the human influence index and precipitation seasonality decline under near-future conditions (**Fig S3, S4**). NDVI, elevation and temperature seasonality were important for the tree-of-heaven model under current and near-future conditions. Frost change frequency also increases in importance for defining tree-of-heaven suitability under near-future climate change conditions in the northeastern US (**Fig S5, Table S10**).

The selection of the mean suitability value between the spotted lanternfly and tree-of-heaven habitat suitability models across all 1km pixels in the northeastern US did not improve the model precision of predicting spotted lanternfly presence (**Table S11**). The selection of the maximum value across the two models did improve model accuracy and decreased uncertainty of estimates across the northeastern U.S. from ROC AUC 97.75% (SD 0.45) to ROC AUC 98.11% (SD 0.18) (**Table S8, S12, Fig S6**). This was the only tree-host suitability model that, when combined with the spotted lanternfly habitat suitability model, improved model performance from the spotted lanternfly distribution model alone (**Table S8, S12**). New York shows a mosaic of increased and decreased suitability from the combined map under near-future climate change conditions while in regions further to the southwest increasing suitability seems to predominate (**Fig S7**). The primary mechanism for the increase in model precision appears to be through increasing suitability estimates for spotted lanternfly in regions that are currently being invaded by the insect through the higher maximum suitability values in the Hudson Valley and Finger Lakes estimated from the tree-of-heaven model (**Fig. 3A, S8**).

**Fig. 3.**
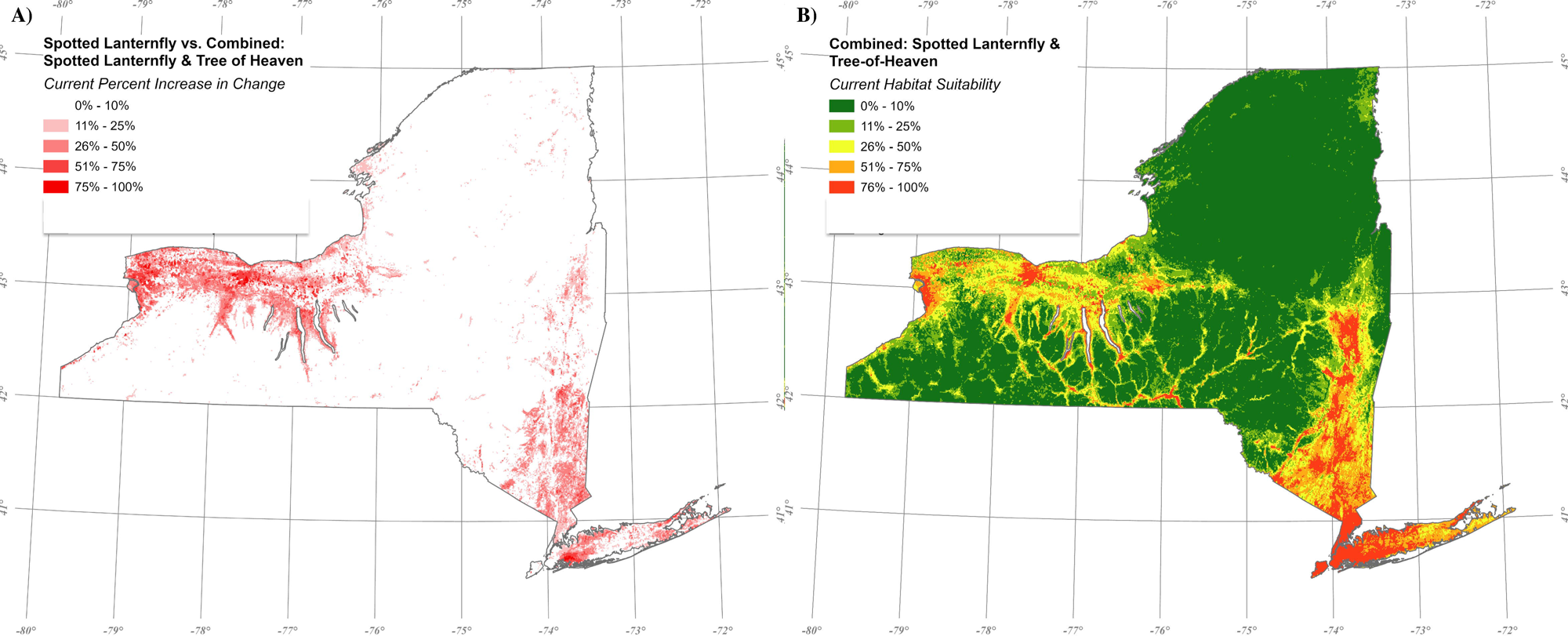
Combined maps of spotted lanternfly and tree-of-heaven suitability in which pixel-wise suitability values were determined by selecting the maximum value between the spotted lanternfly suitability map and the tree-of-heaven suitability map. In A) is the increase in spotted lanternfly suitability resulting from the combination with the tree-of-heaven suitability map, and B) is map illustrating combined tree-of-heaven and spotted lanternfly habitat suitability under current (2011-2040, RCP 7.0) climate conditions in New York state.

Under current conditions, Long Island is the region with the highest suitability for spotted lanternfly with 87% of the region identified to have suitability over 50% (**Fig. 3B**). The Hudson Valley is next with 33% of the region having suitability for spotted lanternfly over 50%, then the Finger Lakes with 15% and finally Lake Erie with only 5%. In the near-future under worsening climatic conditions Long Island decreases the area with high or very high suitability by 5%, while the Finger Lakes, Hudson Valley and Lake Erie remain roughly the same. This could be because Long Island will experience greater precipitation seasonality under near-future climate change conditions while the other three regions will experience declining precipitation seasonality (**Table S13**).

### Vineyard Risk Assessment

Vineyards in the Lake Erie region had the highest vineyard density with an additive risk factor of 7% being applied to the combined spotted lanternfly tree-of-heaven suitability map. Vineyard density in the Finger Lakes an average of 4% additive risk, 7% in Lake Erie, in Long Island 2% and in the Hudson Valley < 1% was added to the combined spotted lanternfly suitability (**Fig. 4A**). Currently, the Long Island region has the highest risk burden with 98% of its vineyards having on-point spotted lanternfly suitability above 50%, categorized as high or very high risk for spotted lanternfly (**Table 1**, **Fig. 4B**). In the Finger Lakes region, 57% of vineyards are at high or very high risk and Hudson Valley vineyards have 38%. Vineyards in the Lake Erie region are dominated by lower risk vineyards with only 12% at high or very high risk despite the added risk due to high vineyard density (**Fig. 4A**, **Table 1**). In the near-future, vineyards at very high risk of spotted lanternfly (suitability > 75%) will increase in the Finger Lakes by 7% and in the Hudson Valley by 2%, however in Long Island vineyards at very high risk will decline from 83% to 61%. Very high-risk vineyards in Lake Erie will remain stable at less than 1% (**Table 1, Fig S9, S10, S11**).

**Fig. 4:**
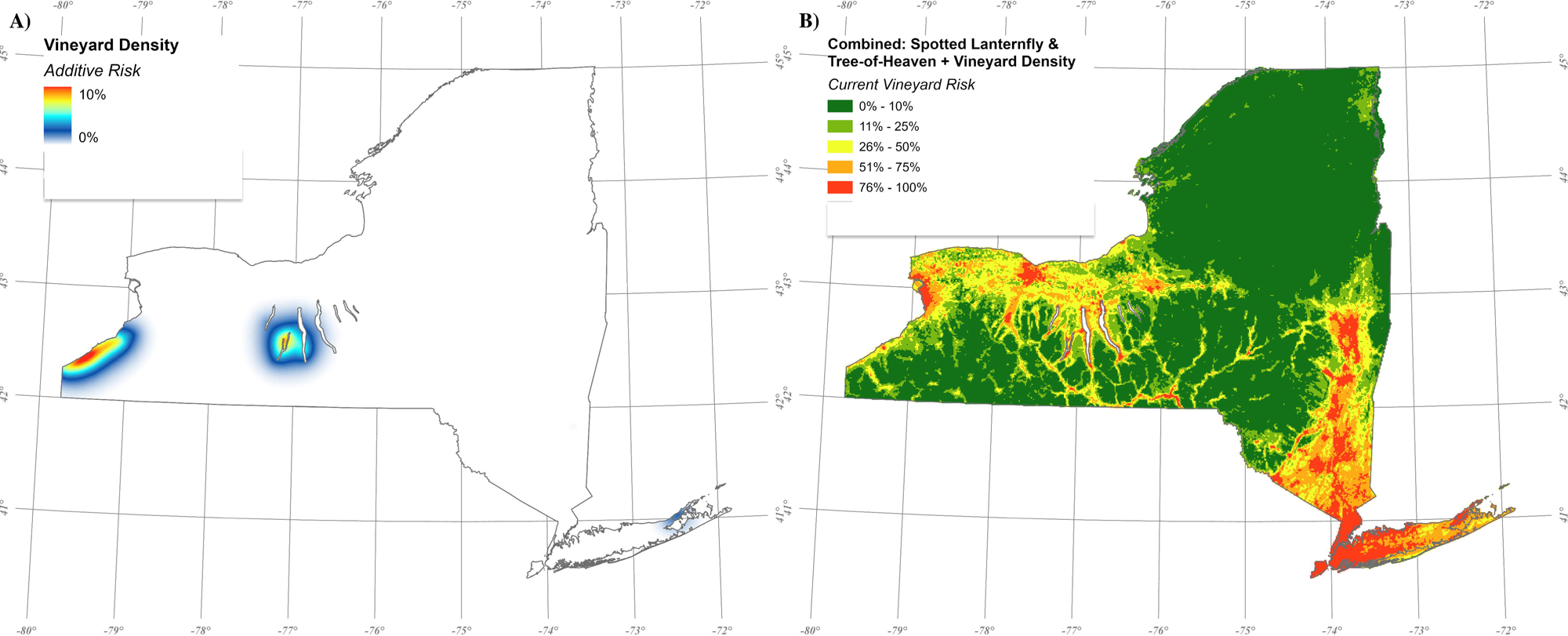
A map illustrating vineyard risk under the current conditions in New York state (RCP 7.0, 2011-2040). In A) the vineyard density additive risk factor is shown in which an adaptive kernel density surface was calculated based on the distance of the nearest 10 neighboring vineyards and using a base influence of 20km and resolution of 1km with maximum additive risk of 10%. In B) the vineyard density surface is added to the combined spotted lanternfly tree-of-heaven suitability map creating a vineyard risk map for spotted lanternfly.

**Table 1:** Percent of vineyards at risk New York state under both the current (SSP 370, RCP 7.0; 2011-2040) and near-future (SSP 585, RCP 8.5; 2041-2070) climate change scenarios; focusing exclusively on vineyard locations in four grape-growing regions: Lake Erie (N=3,700), Finger Lakes (N=2,280), Hudson Valley (N=118), and Long Island (N=381). Vineyards with 0% to 25% risk were categorized low; 25% to 50% risk were categorized moderate; 50% to 75% risk were categorized high; and 75% to 100% risk were categorized very high.

| Current |  |  |  |  |  |
| --- | --- | --- | --- | --- | --- |
| *Risk* | *Risk Category* | *Lake Erie* | *Finger Lakes* | *Hudson Valley* | *Long Island* |
| 0% – 25% | Low | 28% | 13% | 22% | 0% |
| 25% – 50% | Moderate | 59% | 30% | 40% | 2% |
| 50% – 75% | High | 12% | 41% | 29% | 16% |
| 75% – 100% | Very High | 0.4% | 16% | 9% | 83% |
| Future |  |  |  |  |  |
| 0% – 25% | Low | 44% | 9% | 19% | 0% |
| 25% – 50% | Moderate | 43% | 30% | 38% | 0.5% |
| 50% – 75% | High | 13% | 39% | 31% | 38% |
| 75% – 100% | Very High | 0.3% | 22% | 11% | 61% |

## Discussion

The invasive spotted lanternfly is both an economic threat and useful system for modeling invasive species dynamics and how climate change alters suitability. Human influence and near-surface relative humidity are the most important variables predicting habitat suitability for spotted lanternfly under both current and near-future climate conditions. Creating a hybrid map, in which for each pixel across New York state the maximum suitability was selected between the spotted lanternfly suitability map and the tree-of-heaven suitability map, model precision improved in selecting pixels in which spotted lanternfly occurrence was documented. No other alternate tree-hosts aside from tree-of-heaven improved model fit for estimating spotted lanternfly occurrence.

When an additive risk factor for vineyard density was added to the model that could increase risk by a maximum of 10% in high density locations, mean vineyard risk was increase by 7% in the Lake Erie Region and 4% in the Finger Lakes. These regions have higher vineyard densities than either the Hudson Valley or Long Island.

This study demonstrates the value of integrating extensive observational datasets and the distribution of important established plant-hosts with traditional climatic variables to assess the establishment potential of an invasive generalist insect. We show the value of this approach through the increased model estimation of spotted lanternfly occupancy, demonstrating the dynamic interaction between the potential and realized niche of this currently invading species. The addition of vineyard density represents another important methodological approach to quantify added risk of invasive species for important agricultural crops.

The human influence index and near-surface relative humidity emerged as dominant predictors of spotted lanternfly habitat suitability. Human influence index ranked as the most influential variable under both current and future scenarios, which is consistent with previous studies emphasizing that spotted lanternfly spread is strongly dependent on anthropogenic movement and repeated introductions rather than by flight-driven dispersal alone (Ladin et al. 2023b; Elsensohn et al. 2024; Strömbom et al. 2024). Near-surface relative humidity was one of the most influential variables for modeling spotted lanternfly suitability under both the current and future climatic conditions. Higher relative humidity levels reduce spotted lanternfly desiccation and increase egg hatch rate success (Keena 2024; Liu 2022). Currently, the optimal range for spotted lanternfly suitability is between 71% and 84% humidity, increasing along with the projected increase in this variable across New York state in the near-future. Humidity is often overlooked when modeling ectotherm habitat suitability but is arguably one of the most important variables defining the ability of ectotherms to buffer climatic extremes (Brown et al. 2023). Despite its absence in the CHELSA variables using CMIP6, clearly even the estimate used from CMIP4, is still a justified inclusion given the importance of this variable identified by the Random Forest analysis.

New York state is characterized by microclimatic variation with minimum annual extreme temperatures as low as -35°C, making it an important region to help define the cold hardiness limits of spotted lanternfly (Gómez-Marco and Hoddle 2022; Turbelin et al. 2025). Climate change is resulting in increased precipitation, warmer temperatures, and longer growing seasons in New York state, thereby contributing to an environment that favors both grape development and pest survival (Karger et al. 2023). The state’s average temperatures have already increased by approximately 1.7°C since 1970, and projections suggest an additional rise of up to 1.7°C by 2080, particularly in northern regions (Seggos 2021). This warming trend is expected to result in milder winters with fewer days below freezing and reduced snow cover, allowing for a potential range expansion for spotted lanternfly and important hosts like tree-of heaven (Clark et al. 2014). Climate change predictions for New York indicate that while average temperatures are expected to rise, the changes in isothermality are projected to be minimal. This suggests that the relationship between daily and annual temperature variations remain stable, even as overall temperatures increase. The increasing importance of frost change frequency in defining habitat suitability for spotted lanternfly in the future was unexpected. Though freeze-tolerant ectotherms overwintering above ground can experience less physiologically stressful conditions (Voituron et al. 2002; Irwin and Lee 2003), spotted lanternfly eggs are susceptible to damage from repeated freeze-thaw cycles, which can compromise their viability (Irwin and Lee 2003). Frost change frequency may define an important characteristic limiting overwintering success at the colder limits of spotted lanternfly suitability under worsening climate change conditions.

Spotted lanternfly management is inherently challenging because populations frequently disperse among forested, residential, and agricultural landscapes, allowing reinvasion of agricultural fields from surrounding areas (Urban and Leach 2023). Insecticide applications have demonstrated short-term efficacy in suppressing spotted lanternfly populations (Park et al. 2009; Clark et al. 2014; H. Leach et al. 2019). Insecticides may also negatively affect nontarget organisms, including natural enemies and pollinators, thereby limiting their compatibility with integrated pest management strategies (Elmquist et al. 2023). Additionally, these treatments are costly and require repeated applications due to continual reinfestation from adjacent areas that provide alternate plant hosts. Efforts to remove tree-of-heaven have been successful, but they are also resource and labor intensive (Young et al. 2020). The findings in this study emphasize the importance of taking a sub-regional modeling approach to risk assessment. The variation that our models identify at a 1 km gives policymakers a tool to strategically deploy limited resources to targeted locations to mitigate potential negative economic impacts of spotted lanternfly on the New York grape industry.

Sub-regional risk assessments also enable more efficient economic decision-making by stakeholders, allowing them to balance short-term intervention costs with the longer-term benefits of preserving grapevine health and productivity (Chapman et al. 2019). By forecasting potential hotspots under a high-emissions climate change scenario (2041-2070, RCP 8.5), growers can prioritize alternative management practices in regions projected to face higher risks (Harper et al. 2019). Climate change forecasting extends the utility of this model under predicted future climate scenarios in the state of New York.

Though our models provide valuable insights, several limitations should be acknowledged. Under-sampling of both tree-of-heaven and spotted lanternfly in Northern New York and oversampling in New York City may have influenced specific distribution predictions, however strong model predictive power temper these concerns. Future research should concentrate on enhancing sampling methodologies, incorporating finer high-resolution climate data, and investigating the impact of biotic factors, like competition and predation, on species distributions. Another important avenue for exploration involves vineyard-specific risks, since management strategies may further shift vulnerability. This analysis demonstrates the value of combining the maximum suitability between a tree-of-heaven and spotted lanternfly to better estimate overall habitat suitability for the spotted lanternfly in New York state. Though Booth et al. (2025) suggest that other tree species common in New York including walnut, maple (*Acer* spp. L.), and black walnut (*Juglans nigra* L.) are additional tree hosts, we found they did not improve predictive accuracy of estimating spotted lanternfly occurrence. It is still possible, however, that they may bolster populations in marginally suitable conditions.

The overarching goal of this study was to quantify the risk of spotted lanternfly to New York viticulture now and in the near-future by combining vineyard density and tree-of-heaven suitability with ∼16,000 spotted lanternfly observations across the northeastern U.S. We also aimed to identify the probable patterns of future range expansion and contraction under high emissions climate change scenario. Our results indicate that Long Island and the southwestern region of the Finger Lakes are at highest risk of spotted lanternfly establishment and are vineyards there are most likely to experience economic damage. Since range expansion is constrained by many climatic variables which are often expected to shift in opposing directions, this results in a mottled pattern of increased and decreased spotted lanternfly suitability across New York State in the near-future, despite warming temperatures. It is also important to clarify that high suitability for spotted lanternfly does not guarantee invasion. This vineyard risk assessments highlights potential hotspots where spotted lanternfly is more likely to establish and cause economic damage to a vulnerable industry.

As climate change alters our natural environment, implementing proactive pest management strategies, like targeted monitoring, early intervention and the removal of important host plants, becomes essential. The models presented in this study provide valuable insights that can assist stakeholders, policymakers, and growers in making informed decisions about resource allocation, regional monitoring efforts and tree-of-heaven removal. The integration of climate change scenarios quantifies the longer-term local risks by highlighting key variables driving spotted lanternfly range expansion and contraction. Increased temperatures and precipitation patterns can facilitate range expansion for the spotted lanternfly, but extreme temperatures beyond suitability limits in the Hudson Valley and Long Island could potentially put stress on populations. Habitat suitability parameters from the combined spotted lanternfly and tree-of-heaven suitability model suggests that climate change has the potential to contribute to both the expansion and contraction of spotted lanternfly range. This emphasizes the need for targeted, sub-region-specific management strategies. An interactive tool has been developed from the output of the vineyard risk quantified in this assessment. This web-based tool demonstrates the local quantified spotted lanternfly vineyard risk at the 1km scale in New York, and is hosted by the Network for Environmental Weather Applications and Cornell Integrated Pest Management (Emery and Ghashghai 2025). From both environmental and economic perspectives, this tool can be used as a public-facing resource to inform the allocation of local and regional management resources, with the aim of improving the resilience of New York grape production.

## Acknowledgements

We would like to thank the New York State Department of Agriculture and Markets for providing essential datasets for our models. We would like to thank members of Cornell Integrated Pest Management and Cornell Cooperative Extension who provided feedback on interpretation of these findings. We would like to thank Dr. Soheil Boroushaki at California State University, Northridge for his consultation on spatial analyses. This work is supported by the Research Capacity Fund (Hatch), project award number 7009069, from the U.S. Department of Agriculture’s National Institute for Food and Agriculture. This work is also supported by a Cornell University CALS Moonshot Seed Grant (11-24), the New York Wine and Grape Foundation (68141) and NASA Acres (80NSSC23M0034).

