## Supplementary Materials, Supplementary Methods, Table S, Figure S for "High regional variation of spotted lanternfly suitability in New York wine-growing regions signals uneven vineyard risk"

*Data sources*

**1.1** *Occurrence Data Acquisition*

Georeferenced records for five focal taxa were queried from GBIF: *Lycorma delicatula* (White, 1845) (spotted lanternfly; SLF), *Ailanthus altissima* (Mill.) Swingle (tree of heaven), *Juglans nigra* L. (black walnut), *Acer rubrum* L. (red maple), *Acer saccharum* Marshall (sugar maple). Queries were spatially constrained to a defined study extent (*xmin*=-84.8202, *ymin*=36.54031, *xmax*=-66.94983, *ymax*=47.45972) and restricted to records with geographic coordinates. Automated record-level tests were applied to identify and remove potentially problematic coordinates, following the framework implemented by Zizka *et al.* (2019). Records were retained only if coordinate uncertainty was ≤5 km and the basis of record was classified as human observation. Because some biases cannot be detected at the individual record level, dataset-level tests for degree-minute conversion and decimal rounding bias were applied to identify systematic spatial precision errors (Zizka et al., 2019). Cleaning was conducted separately for each species, and only records passing all tests were retained for analysis. For the spotted lanternfly Global Biodiversity Information Facility (GBIF) dataset, records from New York state were removed to avoid duplication when merging with the New York State Department of Agriculture and Markets (NYSDAM) dataset, resulting in a single, complete cleaned dataset.

*Climatic variables*

**2.1** *Downscaling Near-Surface Relative Humidity*

The near-surface relative humidity dataset was statistically downscaled from 1 km spatial resolution to 30 m using universal kriging, with 30 arc second U.S. Geological Survey GMTED2010 elevation as the covariate. Kriging was performed using the *gstat* package (version 2.1-4) (Pebesma, 2004), as it allowed for systematic cross-validation and explicit quantification of accuracy, bias, and interannual stability across variables and years. Similar to Alves et al. (2025), the degree of localization was restricted to the closest 40 training cells for each targeted cell. Cellwise predictions and associated kriging variances were rasterized back to the fine grid to produce monthly downscaled prediction surfaces and corresponding uncertainty (standard error) rasters, which were subsequently cropped and masked to the study boundary and exported as GeoTIFF files.

Model performance was evaluated on a per-day basis using leave-one-out cross-validation (LOO-CV) at the grid-based training locations. For each day, observed values were sequentially withheld and predicted using the same universal kriging specification, variogram model, and neighborhood structure. Cross-validated predictions were summarized (**Table S2**) using root mean squared error (RMSE) to measure the mean performance calculated across all daily raster outputs under both current (2011-2040; RCP 7.0) and future (2041-2070; RCP 8.5) scenarios.

*Geographic variables*

**3.1** *Vegetation phenology and productivity*

The Normalized Difference Vegetation Index (NDVI) dataset was acquired through the Harmonized Landsat Sentinel-2 (HLS) project. The Operational Land Imager (OLI) is located on the joint NASA/USGS Landsat 8 and Landsat 9 satellites, whereas the Multi-Spectral Instrument (MSI) is mounted on Europe's Copernicus Sentinel-2A, Sentinel-2B, and Sentinel-2C satellites. The combined measurement enables global observations of the land every two to three days at 30 m spatial resolution. Using Google Earth Engine (Gorelick *et al.*, 2017, https://code.earthengine.google.com/), we extracted the red (band 4) and near-infrared (band 8) bands between August 1, 2024, and November 1, 2024, and calculated the NDVI before exporting it to Google Drive; with values ranging between -1 and 1 (**Table S5**). The extent included New York, Pennsylvania, Virginia, West Virginia, Ohio, Maryland, New Jersey, Delaware, Connecticut, Delaware, Massachusetts, New Hampshire, Rhode Island, Vermont, and Maine.

**3.2** *Human influence*

The Human Influence Index compiles information from remote sensing data that quantifies human-dominated land cover, population density, power utility infrastructure, agricultural lands, roads, railways, and navigable waterways on a global scale (Venter et al., 2024). The human footprint is then calculated as an index of total human pressure from those combined indices of land-use pressure, ranging from low to very high (**Table S6**).

*Spotted lanternfly habitat suitability*

**4.1** *Data Preprocessing*

The preprocessing stage involves the preparation and refinement of spatial occurrence data to ensure consistency and accuracy in downstream analyses. Initially, remotely sensed raster images are downloaded and masked to the study area extent. The extent included New York, Pennsylvania, Virginia, West Virginia, Ohio, Maryland, New Jersey, Delaware, Connecticut, Delaware, Massachusetts, New Hampshire, Rhode Island, Vermont, and Maine. All raster datasets were reprojected to a common Coordinate Reference System (CRS=4326) to ensure spatial alignment and then resampled to a common resolution (1 km) using a bilinear approach. Subsequently, all raster layers are then stacked into a multi-band dataset to standardize input variables.

Occurrence data was uploaded, and missing values were identified and removed to maintain consistency. A spatial points object was created from a structured data frame to establish a geospatial reference. To balance the dataset, pseudo-absence points were generated by randomly sampling an equal number of absences and presences, then encoded with binary classification labels: presence (1) and absence (0). The presences and absences were then filtered out separately based on binary classification labels (0, 1). We combined 70% absences and 70% presences to be used for training and testing our model. Then, we combined the remaining 30% absences and 30% presences into a separate independent dataset to be used for validating our model.

**4.2** *Model Assessment*

The testing-training and validation datasets were then analyzed with the pre-stacked raster dataset of climatic and geographic variables. The validation dataset remained unaltered, while the training-testing dataset underwent internal partitioning using the bootstrap resampling method. Within the training-testing dataset, present and absence data was randomly divided, with 80% allocated for training and 20% for testing, repeated across ten replicates. Spatial information was then extracted from the pre-stacked raster dataset using latitude and longitude coordinates via training-testing dataset.

The training-testing dataset was initially transformed into a spatial format and aligned with a common Coordinate Reference System (CRS = 4326), corresponding to the stacked raster dataset. Using the 'cv-spatial' function, spatial blocks were generated at a 1 km radius, with tenfold partitioning and 100 randomized iterations. The response variable was then utilized to extract relevant explanatory variables via the stacked raster dataset. The 'train_control' function was used to implement cross-validation using the precomputed spatial folds while storing only the final predictions. A Random Forest model was then trained with a tuning length of five and ntree = 200. The variable importance of all explanatory variables was assessed. Model performance was then evaluated for the model using current climatic conditions using our independent validation dataset, generating a confusion matrix; particularly looking at accuracy, sensitivity, specificity, positive prediction value, and negative prediction value.

#### **4.3** Tree *of Heaven Suitability Mapping*

The climatic and geographic variables driving regional differentiation for tree of heaven habitat suitability included isothermality, frost change frequency, temperature seasonality, precipitation seasonality, monthly precipitation of the coldest quarter, elevation, NDVI, and soil texture. The tree of heaven model under the current (2011-2040; RCP 7.0) scenario demonstrated high accuracy (87.76%), sensitivity (86.60%), specificity (88.94%), positive prediction value (88.82%), and negative prediction value (86.74%).

According to the Mean Decrease Gini, the three most influential explanatory variables for modeling tree of heaven between both current and future scenarios, from highest to lowest importance, were NDVI, elevation, and temperature seasonality (**Figure S2**). Interestingly, frost change frequency increases in importance under the future scenario, with a Mean Decrease Gini value of 478 under current conditions that increases to 737. The highest suitability for tree of heaven is characterized by dense to very dense vegetation, elevation between 185 m and 1,665 m under both current and future conditions, and temperature seasonality above 937°C under current conditions and above 897°C under future conditions (**Table S7**).

#### **4.4** *Black Walnut Suitability Mapping*

The climatic and geographic variables driving regional differentiation for black walnut habitat suitability included isothermality, frost change frequency, temperature seasonality, precipitation seasonality, monthly precipitation of the coldest quarter, elevation, NDVI, and soil texture. The black walnut model under the current (2011-2040; RCP 7.0) scenario demonstrated high accuracy

(82.82%), sensitivity (82.18%), specificity (83.47%), positive prediction value (83.35%), and negative prediction value (82.31%).

According to the Mean Decrease Gini, the three most influential explanatory variables for modeling black walnut between both current and future scenarios, from highest to lowest importance, were elevation, NDVI, and temperature seasonality (**Figure S2**). Interestingly, frost change frequency increases in importance under the future scenario, with a Mean Decrease Gini value of 586 under current conditions that increases to 767. The highest suitability for black walnut is characterized by dense to very dense vegetation, elevation between 280 m and 1,581 m under current conditions and between 225 m and 1393 m under future conditions, and temperature seasonality above 931°C under current conditions and above 897°C under future conditions (**Table S7**).

**4.5** *Red Maple Suitability Mapping*

The climatic and geographic variables driving regional differentiation for red maple habitat suitability included isothermality, frost change frequency, temperature seasonality, precipitation seasonality, monthly precipitation of the coldest quarter, elevation, NDVI, and soil texture. The red maple model under the current (2011-2040; RCP 7.0) scenario demonstrated high accuracy (81.12%), sensitivity (82.00%), specificity (80.24%), positive prediction value (80.59%), and negative prediction value (81.67%).

According to the Mean Decrease Gini, the three most influential explanatory variables for modeling red maple between both current and future scenarios, from highest to lowest importance, were elevation, isothermality, and temperature seasonality (**Table S2**). Interestingly, the mean monthly precipitation of the coldest quarter increases in importance under the future scenario, with a Mean Decrease Gini value of 643 under current conditions that increases to 748. Precipitation seasonality, however, decreases in importance under the future scenario, with a Mean Decrease Gini value of 856 under current conditions that decreases to 606. The highest suitability for red maple is characterized by an elevation between 159 m and 1,581 m under both current and future conditions, isothermality between 13% and 33% under current conditions and between 13% and 36% under future conditions, and temperature seasonality above 745°C under current conditions and above 734°C under future conditions (**Table S7**).

**4.6** *Sugar Maple Suitability Mapping*

The climatic and geographic variables driving regional differentiation for sugar maple habitat suitability included isothermality, frost change frequency, temperature seasonality, precipitation seasonality, monthly precipitation of the coldest quarter, elevation, NDVI, and soil texture. The sugar maple model under the current (2011-2040; RCP 7.0) scenario demonstrated high accuracy (83.20%), sensitivity (84.64%), specificity (81.76%), positive prediction value (82.25%), and negative prediction value (84.21%).

According to the Mean Decrease Gini, the three most influential explanatory variables for modeling sugar maple between both current and future scenarios, from highest to lowest importance, were isothermality, temperature seasonality, and frost change frequency (**Table S2**). Interestingly, the isothermality increases in importance under the future scenario, with a Mean Decrease Gini value of 820 under current conditions that increases to 954. Frost change frequency, however, decreases in importance under the future scenario, with a Mean Decrease Gini value of 882 under current conditions that decreases to 755. The highest suitability for sugar maple is characterized by isothermality between 12% and 35% under current conditions and between 13% and 36% under future conditions, temperature seasonality between 617°C and 1,022°C under current conditions and above 582°C under future conditions, and frost change frequency with a maximum of 81 events under current conditions and a maximum of 143 events under future conditions (**Table S7**).

### Supplementary Tables and Figures

**Table S1:** The full range of each predictor variables under both current (2011-2040; RCP 7.0) and near-future (2041-2070; RCP 8.5) climate change scenarios, when future predictions are available, in northeastern United States. States included New York, Pennsylvania, Virginia, West Virginia, Ohio, Maryland, New Jersey, Connecticut, Massachusetts, New Hampshire, Rhode Island, Vermont, and Maine. (+) indicates an increase relative to current conditions, while the (-) indicates a decrease.

| Explanatory Variable | Resolution | Unit | Current Range | Future Range | Change |
| --- | --- | --- | --- | --- | --- |
| Elevation | 30 m | m | 0 – 1389 | NA | |
| Topographic Position Index | 30 m | | -195.5 – 190.5 | NA | |
| Soil Texture | 250 m | | 1 – 11 | NA | |
| Tree Canopy Cover | 30 m | % | 0 – 100 | NA | |
| Normalized Difference Vegetation Index | 30 m | | -1.0 – 1.0 | NA | |
| Human Influence Index | 1 km | | 1 – 50 | NA | |
| Near-Surface Relative Humidity | 30 m | % | 70 – 81 | 69 – 77 | - |
| Isothermality | 1 km | % | 15.3 – 33.9 | 15.2 – 36.5 | + |
| Frost Change Frequency | 1 km | count | 0 – 140 | 0 – 142 | + |
| Temperature Seasonality | 1 km | °C | 616.1 – 1034.0 | 582.9 – 988.2 | - |
| Precipitation Seasonality | 1 km | kg $m^{-2}$ | 112 – 268 | 95 – 221 | - |
| Mean Monthly Precipitation of the Coldest Quarter | 1 km | kg $m^{-2}$ $month^{-1}$ | 1761 – 4317 | 1987 – 5074 | + |

**Table S2:** A table summarizing cross-validation performance metrics for the downscaling of near-surface relative humidity; focusing on root mean squared error (RMSE), absolute error (MAE), bias, and the coefficient of determination ($R^2$). Reported values represent mean performance metrics calculated across all daily raster outputs under both current (2011-2040; RCP 7.0) and future (2041-2070; RCP 8.5) scenarios.

| Near-Surface Relative Humidity | | |
| --- | --- | --- |
| | *Current* | *Future* |
| RMSE | 1.5225 | 1.3510 |
| MAE | 1.1906 | 1.0274 |
| Bias | -0.0087 | -0.0580 |
| $R^2$ | 0.1730 | 0.0638 |

**Table S3:** Classes represented in the topographic position index raster dataset in northeastern United States. States included New York, Pennsylvania, Virginia, West Virginia, Ohio, Maryland, New Jersey, Connecticut, Massachusetts, New Hampshire, Rhode Island, Vermont, and Maine.

| Topographic Classes | |
|---|---|
| *Value* | *Topographic Type* |
| Positive (>0) | Ridges, Crests, Hilltops |
| Zero | Flat Areas, Mid-Slopes |
| Negative (<0) | Valleys, Depressions, Drainage Lines |

**Table S4:** Classes represented in the soil texture raster dataset in northeastern United States. States included New York, Pennsylvania, Virginia, West Virginia, Ohio, Maryland, New Jersey, Connecticut, Massachusetts, New Hampshire, Rhode Island, Vermont, and Maine.

| Soil Texture Classes | |
|---|---|
| *Value* | *Soil Type* |
| 1 – 3 | Clay and Loam |
| 4 – 5 | Clay and Sand |
| 6 – 8 | Sand, Clay, and Loam |
| 9 – 11 | Silt, Clay, and Loam |

**Table S5:** Classes represented in the normalized difference vegetation index raster dataset in northeastern United States. States included New York, Pennsylvania, Virginia, West Virginia, Ohio, Maryland, New Jersey, Connecticut, Massachusetts, New Hampshire, Rhode Island, Vermont, and Maine.

| NDVI Classes | |
|---|---|
| *Value* | *Vegetation Type* |
| -1.0 – 0.0 | No Vegetation (e.g., water, snow, clouds) |
| 0.0 – 0.2 | Bare Soil, Rock, Urban Surfaces |
| 0.2 – 0.4 | Sparse Vegetation (e.g., grasslands) |
| 0.4 – 0.6 | Moderate Vegetation (e.g., shrublands) |
| 0.6 – 0.8 | Dense, Healthy Vegetation (e.g., forest canopy) |
| 0.8 – 1.0 | Very Dense Canopy (e.g., high leaf area index) |

**Table S6:** Classes represented in the human influence index raster dataset in northeastern United States. States included New York, Pennsylvania, Virginia, West Virginia, Ohio, Maryland, New Jersey, Connecticut, Massachusetts, New Hampshire, Rhode Island, Vermont, and Maine.

| Human Influence Index Classes | |
|---|---|
| *Value* | *Pressure Type* |
| 0 – 12 | Low |
| 13 – 25 | Moderate |
| 26 – 40 | High |
| 41 – 50 | Very High |

**Table S7:** Performance metrics from the spotted lanternfly Random Forest model under the current (SSP 370, RCP 7.0; 2011-2040) and near-future (SSP 585, RCP 8.5; 2041-2070) climate change scenarios in northeastern United States. States included New York, Pennsylvania, Virginia, West Virginia, Ohio, Maryland, New Jersey, Connecticut, Massachusetts, New Hampshire, Rhode Island, Vermont, and Maine. Metrics were determined by cross-validating presence and pseudo-absence spotted lanternfly observations across the original spotted lanternfly Random Forest model outputs. Metrics included accuracy, sensitivity, positive prediction value and negative prediction value.

| Spotted Lanternfly Random Forest Model | |
|---|---|
| Accuracy | 0.9229 |
| Sensitivity | 0.9288 |
| Specificity | 0.9168 |
| Positive Prediction Value | 0.9199 |
| Negative Prediction Value | 0.9260 |

**Table S8**: Performance metrics from evaluating the spotted lanternfly habitat suitability map (without tree-host habitat suitability) under the current (SSP 370, RCP 7.0; 2011-2040) climate change scenario in northeastern United States. States included New York, Pennsylvania, Virginia, West Virginia, Ohio, Maryland, New Jersey, Connecticut, Massachusetts, New Hampshire, Rhode Island, Vermont, and Maine. Metrics were determined by cross-validating presence-only spotted lanternfly observations across the original spotted lanternfly Random Forest model output. Metrics included the receiver operating characteristic area under the curve (ROC AUC) and precision recall area under the curve (PR AUC), along with their corresponding standard deviations.

| Spotted Lanternfly Cross-Validation | |
|---|---|
| ROC AUC | 0.9775 |
| Standard Deviation | 0.0045 |
| PR-AUC | 0.9791 |
| Standard Deviation | 0.0043 |

**Table S9:** The minimum and maximum marginal relationship between individual predictor variables and the predicted probability of spotted lanternfly presence derived from the Random Forest models under both the current (2011-2040; RCP 7.0) and near-future (2041-2070; RCP 8.5) climate change scenarios in northeastern United States (New York, Pennsylvania, Virginia, West Virginia, Ohio, Maryland, New Jersey, Connecticut, Massachusetts, New Hampshire, Rhode Island, Vermont, and Maine).

| Spotted Lanternfly | | | |
|---|---|---|---|
| *Explanatory Variable* | *Unit* | *Current* | *Future* |
| Human Influence Index | | 0 – 22 | 0 – 23 |
| Near-Surface Relative Humidity | % | 73 – 80 | 72 – 76 |
| Temperature Seasonality | °C | 938.3 – 1034.0 | 895.2 – 988.2 |
| Isothermality | % | 20 – 25 | 20 – 27 |
| Tree Canopy Cover | % | 0 – 89 | 57 – 88 |
| Precipitation Seasonality | kg $m^{-2}$ | 84 – 244 | 149 – 210 |
| Frost Change Frequency | count | 68 – 96 | 80 – 139 |
| Mean Monthly Precipitation of the Coldest Quarter | kg $m^{-2}$ $month^{-1}$ | 2334 – 4317 | 2287 – 5074 |
| Topographic Position Index | | -34 – 172 | -43 – 172 |

**Table S10:** The minimum and maximum marginal relationship between individual predictor variables and the predicted probability of tree-of-heaven, black walnut, red maple, and sugar maple presence derived from the Random Forest models under both the current (2011-2040; RCP 7.0) and near-future (2041-2070; RCP 8.5) climate change scenarios in northeastern United States (New York, Pennsylvania, Virginia, West Virginia, Ohio, Maryland, New Jersey, Connecticut, Massachusetts, New Hampshire, Rhode Island, Vermont, and Maine).

| Tree-of-Heaven | | | |
|---|---|---|---|
| *Explanatory Variable* | *Unit* | *Current* | *Future* |
| Elevation | m | 185 – 1665 | 185 – 1665 |
| Frost Change Frequency | count | 75 – 97 | 103 – 136 |
| Isothermality | % | 20.7 – 26.2 | 20.1 – 29.3 |
| Normalized Difference Vegetation Index | | 0.65 – 0.87 | 0.68 – 87 |
| Mean Monthly Precipitation of the Coldest Quarter | kg $m^{-2}$ $month^{-1}$ | 2516 – 5007 | 2618 – 5512 |
| Precipitation Seasonality | kg $m^{-2}$ | 158 – 232 | 99 – 250 |
| Soil Texture | | 0 – 11 | 0 – 11 |
| Temperature Seasonality | °C | 937.5 – 1034.0 | 897.5 – 988.2 |

| | | | |
|---|---|---|---|
| Black Walnut | | | |
| Elevation | m | 280 – 1581 | 225 – 1393 |
| Frost Change Frequency | count | 73 – 122 | 69 – 138 |
| Isothermality | % | 20.5 – 31.6 | 13.4 – 33.2 |
| Normalized Difference Vegetation Index | | 0.72 – 0.88 | 0.71 – 0.88 |
| Mean Monthly Precipitation of the Coldest Quarter | kg $m^{-2}$ $month^{-1}$ | 2595 – 5043 | 2516 – 5610 |
| Precipitation Seasonality | kg $m^{-2}$ | 167 – 265 | 95 – 251 |
| Soil Texture | | 7 – 11 | 0 – 11 |
| Temperature Seasonality | °C | 931.3 – 1034.0 | 897.0 – 988.2 |
| Red Maple | | | |
| *Explanatory Variable* | *Unit* | *Current* | *Future* |
| Elevation | m | 159 – 1581 | 159 – 1581 |
| Frost Change Frequency | count | 0 – 123 | 16 – 145 |
| Isothermality | % | 13.1 – 32.8 | 13.4 – 36.0 |
| Normalized Difference Vegetation Index | | -0.77 – 0.88 | -0.77 – 0.88 |
| Mean Monthly Precipitation of the Coldest Quarter | kg $m^{-2}$ $month^{-1}$ | 1474 – 5040 | 1629 – 3255 |
| Precipitation Seasonality | kg $m^{-2}$ | 99 – 268 | 161 – 257 |
| Soil Texture | | 0 – 11 | 0 – 11 |
| Temperature Seasonality | °C | 745.9 – 1034.0 | 734.1 – 988.2 |
| Sugar Maple | | | |
| Elevation | m | 0 – 1665 | 0 – 1665 |
| Frost Change Frequency | count | 0 – 81 | 0 – 143 |
| Isothermality | % | 12.9 – 34.6 | 13.4 – 36.1 |
| Normalized Difference Vegetation Index | | 0.33 – 0.87 | -0.64 – 0.87 |
| Mean Monthly Precipitation of the Coldest Quarter | kg $m^{-2}$ $month^{-1}$ | 2445 – 5007 | 1641 – 5512 |
| Precipitation Seasonality | kg $m^{-2}$ | 83 – 266 | 82 – 249 |
| Soil Texture | | 0 – 10 | 0 – 10 |
| Temperature Seasonality | °C | 617.3 – 1022.0 | 582.9 – 988.2 |

**Table S11**: Performance metrics from evaluating the integration of tree-host habitat suitability maps with the spotted lanternfly habitat suitability map based on the 'mean' operator under the current (SSP 370, RCP 7.0; 2011-2040) climatic scenario in northeastern United States. States included New York, Pennsylvania, Virginia, West Virginia, Ohio, Maryland, New Jersey, Connecticut, Massachusetts, New Hampshire, Rhode Island, Vermont, and Maine. Metrics were determined by cross-validating presence-only spotted lanternfly observations across each combined suitability map. Metrics included the receiver operating characteristic area under the curve (ROC AUC) and precision recall area under the curve (PR AUC), along with their corresponding standard deviations.

| 'Mean' Operator | | | | | |
|---|---|---|---|---|---|
| | Spotted Lanternfly & All Tree-Hosts | Spotted Lanternfly & Tree of Heaven | Spotted Lanternfly & Black Walnut | Spotted Lanternfly & Red Maple | Spotted Lanternfly & Sugar Maple |
| ROC AUC | 0.9714 | 0.9731 | 0.9647 | 0.9620 | 0.9570 |
| Standard Deviation | 0.0027 | 0.0027 | 0.0040 | 0.0031 | 0.0040 |
| PR-AUC | 0.9670 | 0.9704 | 0.9591 | 0.9575 | 0.9539 |
| Standard Deviation | 0.0049 | 0.0045 | 0.0050 | 0.0065 | 0.0056 |

**Table S12**: Performance metrics from evaluating the integration of tree-host habitat suitability maps with the spotted lanternfly habitat suitability map based on the 'maximum' operator under the current (SSP 370, RCP 7.0; 2011-2040) climatic scenarios in northeastern United States. States included New York, Pennsylvania, Virginia, West Virginia, Ohio, Maryland, New Jersey, Connecticut, Massachusetts, New Hampshire, Rhode Island, Vermont, and Maine. Metrics were determined by cross-validating presence-only spotted lanternfly observations across each combined suitability map. Metrics included the receiver operating characteristic area under the curve (ROC AUC) and precision recall area under the curve (PR AUC), along with their corresponding standard deviations.

| 'Maximum' Operator | | | | | |
|---|---|---|---|---|---|
| | Spotted Lanternfly & All Tree-Hosts | Spotted Lanternfly & Tree of Heaven | Spotted Lanternfly & Black Walnut | Spotted Lanternfly & Red Maple | Spotted Lanternfly & Sugar Maple |
| ROC AUC | 0.9703 | 0.9811 | 0.9684 | 0.9638 | 0.9568 |
| Standard Deviation | 0.0025 | 0.0018 | 0.0041 | 0.0036 | 0.0060 |
| PR-AUC | 0.9715 | 0.9813 | 0.9701 | 0.9678 | 0.9639 |
| Standard Deviation | 0.0025 | 0.0022 | 0.0037 | 0.0031 | 0.0044 |

**Table S13**: Climatic variables under both the current (SSP 370; RCP 7.0; 2011-2040) and near-future (SSP 585, RCP 8.5; 2041-2070) climate change scenarios in New York state, highlighting the directionality in change. Directionality in change was computed using zonal statistics in ArcGIS Pro (v3.3); focusing on four grape-growing regions: Lake Erie, Finger Lakes, Hudson Valley, and Long Island. (+) indicates an increase relative to current conditions, while the (-) indicates a decrease.

| Explanatory Variable | Lake Erie | Finger Lakes | Hudson Valley | Long Island |
|---|---|---|---|---|
| Near-Surface Relative Humidity | + | + | + | + |
| Isothermality | + | + | + | + |
| Frost Change Frequency | - | - | - | - |
| Temperature Seasonality | - | - | - | - |
| Precipitation Seasonality | - | - | - | + |
| Mean Monthly Precipitation of the Coldest Quarter | + | + | + | + |

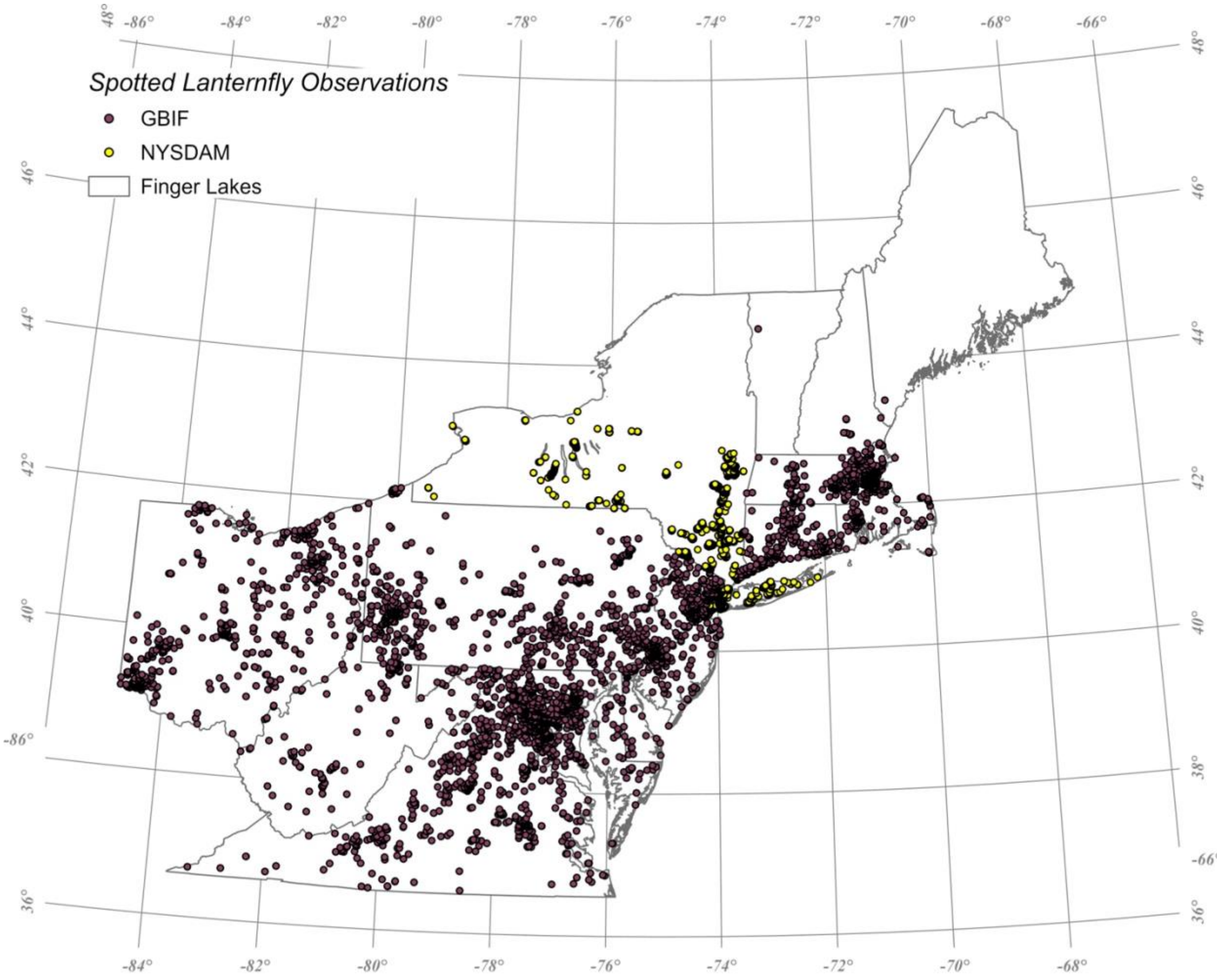

**Figure S1**: Spotted lanternfly occurrences from (yellow) the New York State Department of Agriculture and Markets (NYSDAM; N=10,114) and (burgundy) the Global Biodiversity Information Facility (GBIF; N=5,884) repository databases, resulting in 15,998 total observations in northeastern United States. States included New York, Pennsylvania, Virginia, West Virginia, Ohio, Maryland, New Jersey, Connecticut, Massachusetts, New Hampshire, Rhode Island, Vermont, and Maine.

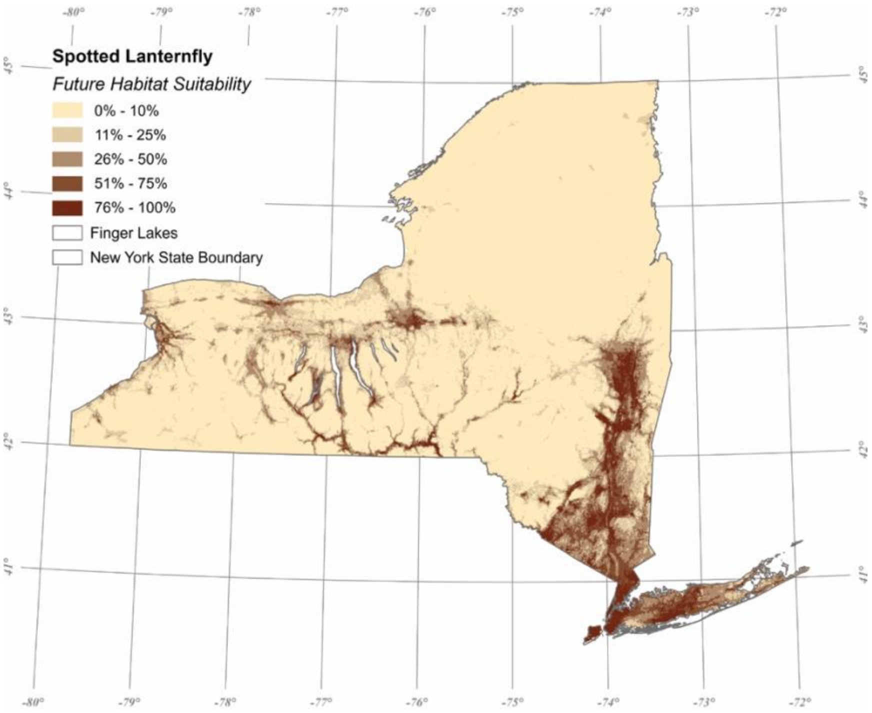

**Figure S2:** A map illustrating the near-future (2041-2070; RCP 8.5) habitat suitability for spotted lanternfly in New York state.

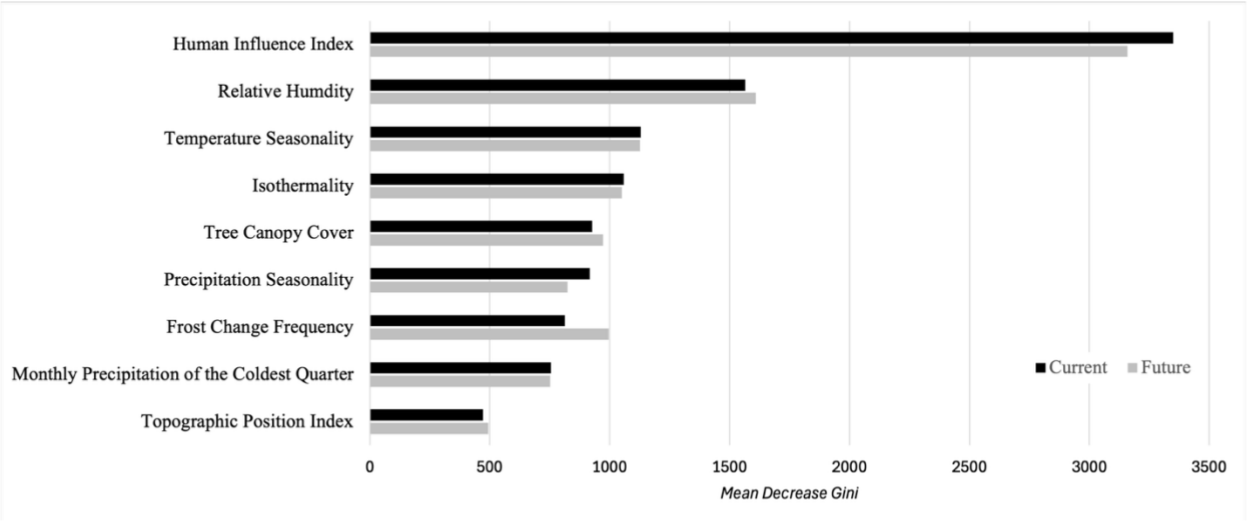

**Figure S3:** Variable Importance based on the Mean Decrease Gini Index for spotted lanternfly Random Forest models under both the (black) current (2011-2040; RCP 7.0) and (grey) near-future (2041-2070; RCP 8.5) climate change scenarios in northeastern United States. States included New York, Pennsylvania, Virginia, West Virginia, Ohio, Maryland, New Jersey, Connecticut, Massachusetts, New Hampshire, Rhode Island, Vermont, and Maine.

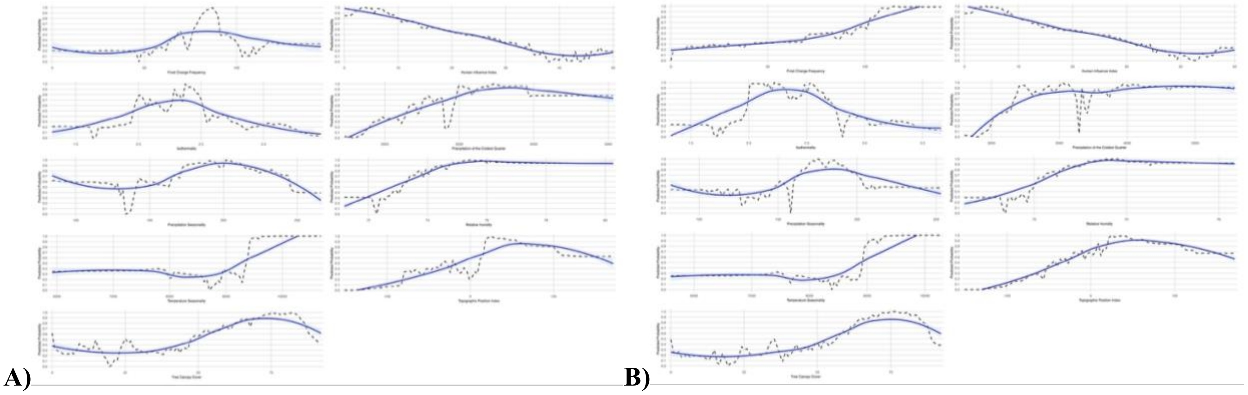

**Fig S4:** Partial dependence plots for the nine predictor variables used in the spotted lanternfly Random Forest model under the A) current (2011-2040; RCP 7.0) and B) the near-future (2041-2070; RCP 8.5) climate change scenario in northeastern United States. The dashed black line in each plot represents the true model response and the blue loess-smoothed curve represents the overall trend. States included New York, Pennsylvania, Virginia, West Virginia, Ohio, Maryland, New Jersey, Connecticut, Massachusetts, New Hampshire, Rhode Island, Vermont, and Maine.

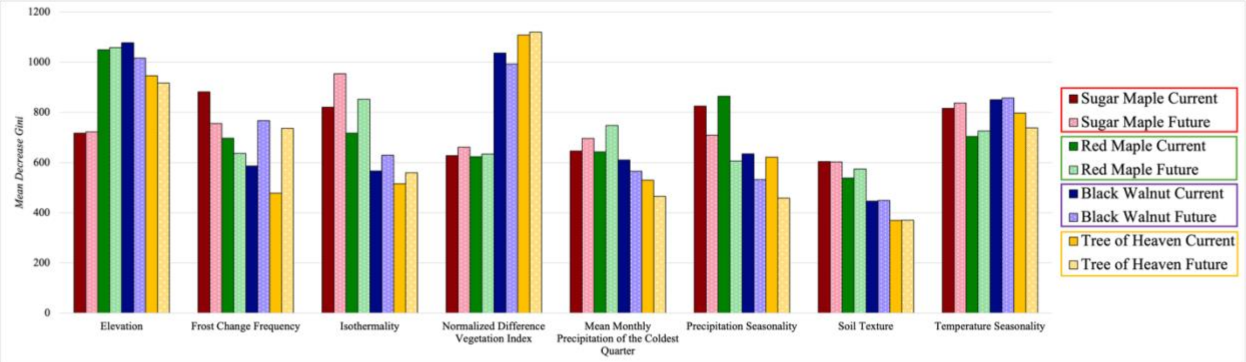

**Figure S5:** Variable Importance based on the Mean Decrease Gini Index for tree of heaven (yellow), black walnut (purple), red maple (green), and sugar maple (red) Random Forest models under both the current (2011-2040; RCP 7.0) and near-future (2041-2070; RCP 8.5) climate change scenarios in northeastern United States. States included New York, Pennsylvania, Virginia, West Virginia, Ohio, Maryland, New Jersey, Connecticut, Massachusetts, New Hampshire, Rhode Island, Vermont, and Maine.

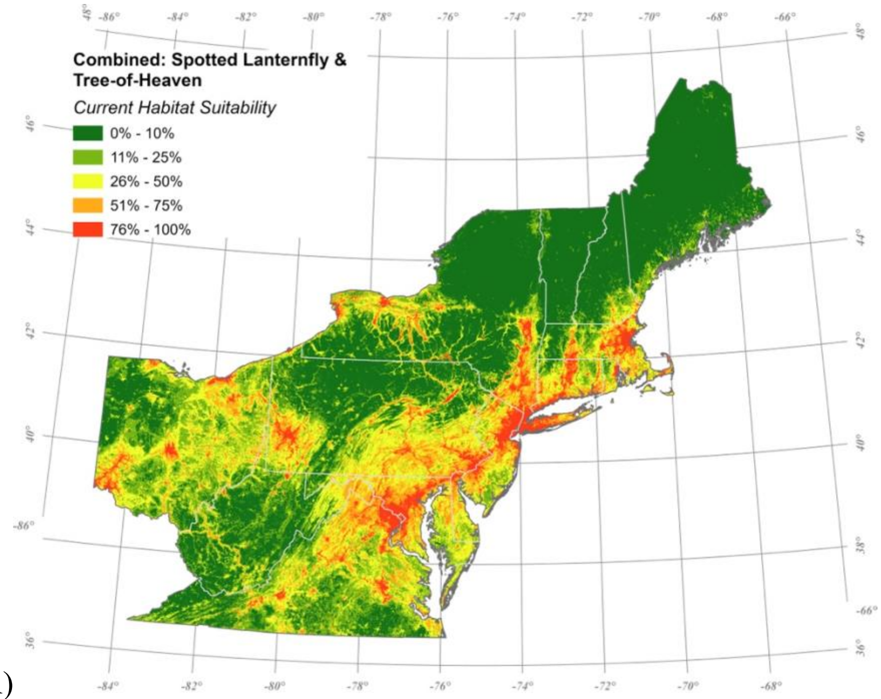

A)

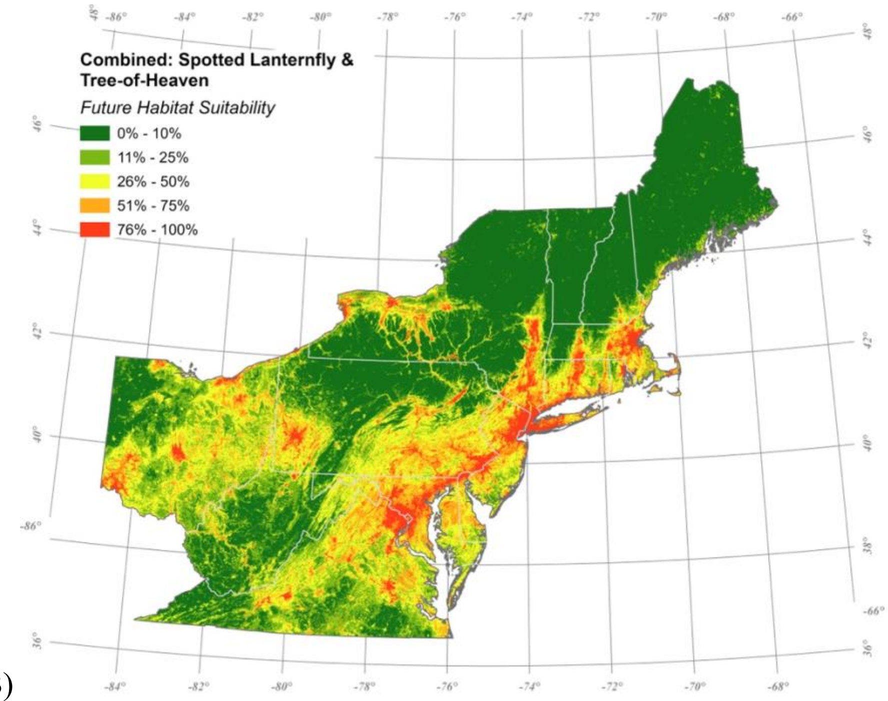

B)

**Figure S6**: A map illustrating the combined habitat suitability map of spotted lanternfly and tree-of-heaven suitability in A) under the current conditions in (2011-2040; RCP 7.0) and in B) under the near-future (2041-2070; RCP 8.5) climate change scenario in northeastern United States. Pixel-wise values are determined by the highest (maximum) suitability between species. All modeled states are shown including New York, Pennsylvania, Virginia, West Virginia, Ohio, Maryland, New Jersey, Connecticut, Massachusetts, New Hampshire, Rhode Island, Vermont, and Maine.

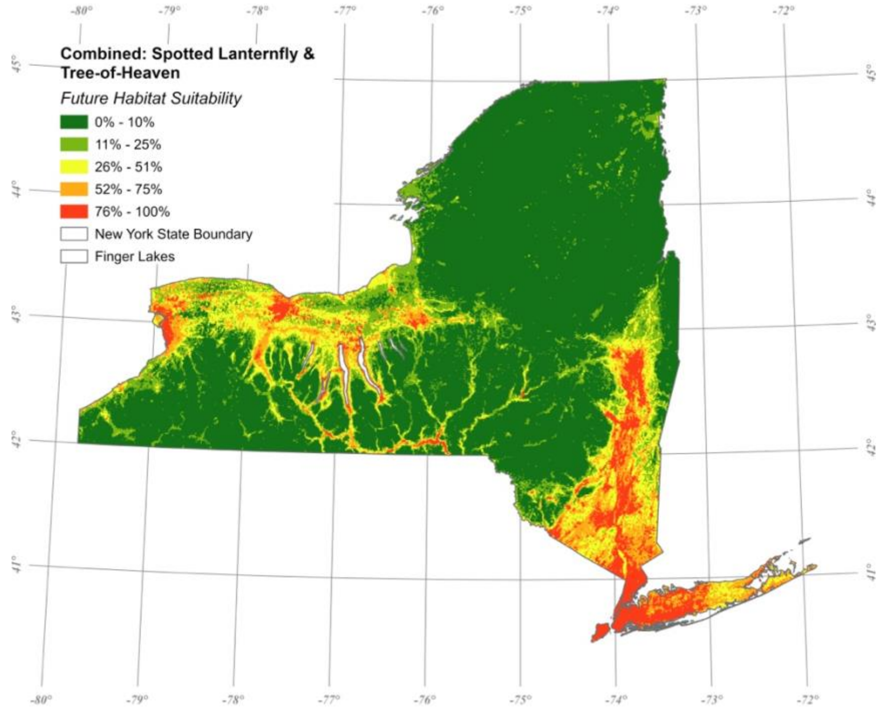

**Figure S7**: A map illustrating the combined habitat suitability map of spotted lanternfly and tree-of-heaven suitability under the near-future (2041-2070; RCP 8.5) climate change scenario in New York state. Pixel-wise values were determined by the maximum suitability between species.

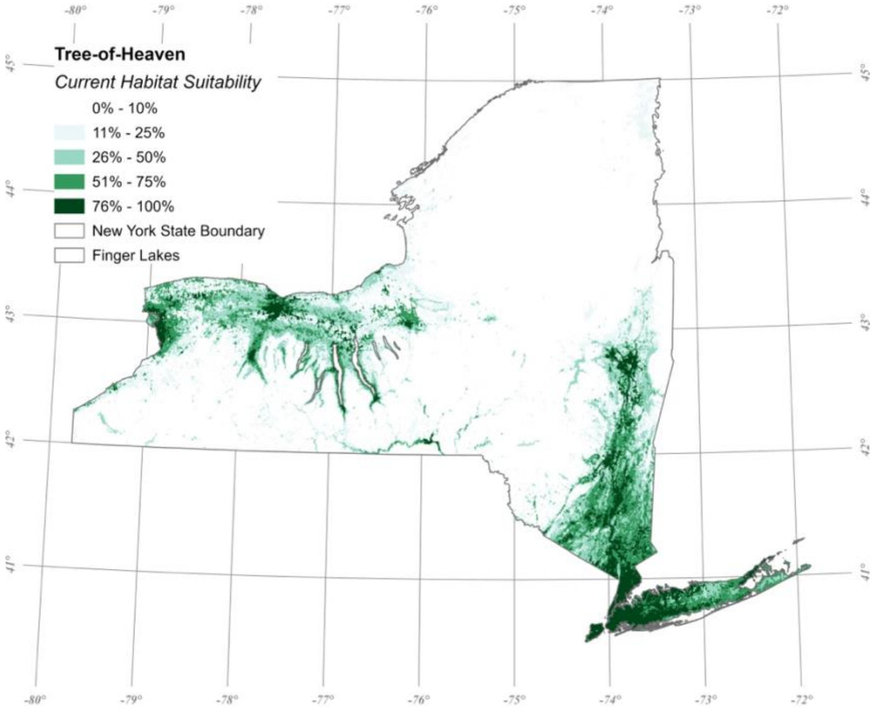

A)

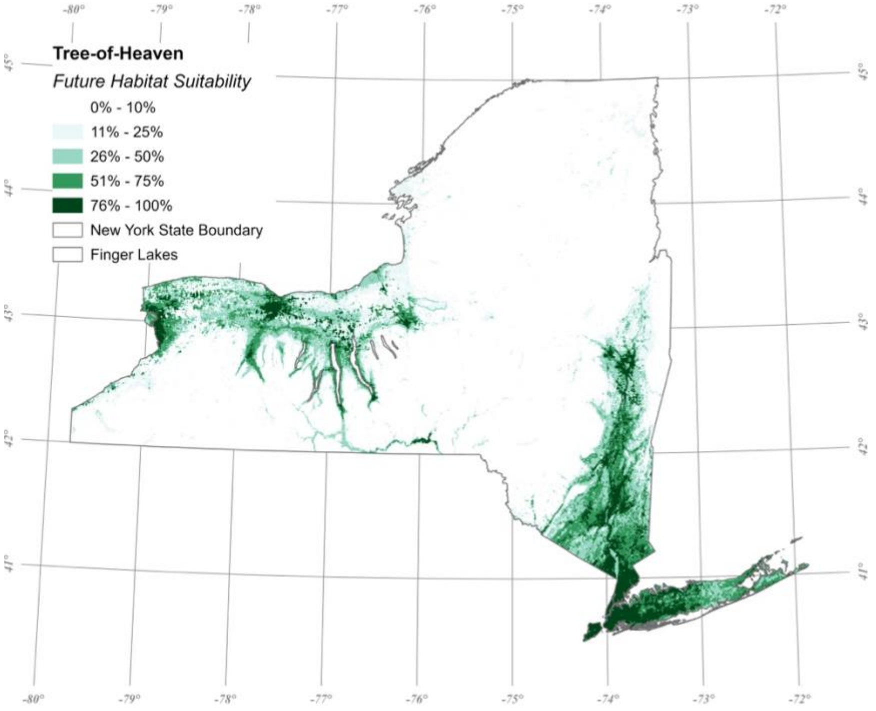

B)

**Figure S8:** A map illustrating the habitat suitability for tree-of-heaven under A) current (2011-2040; RCP 7.0), and B) near-future (2041-2070; RCP 8.5) climate change scenarios in New York state.

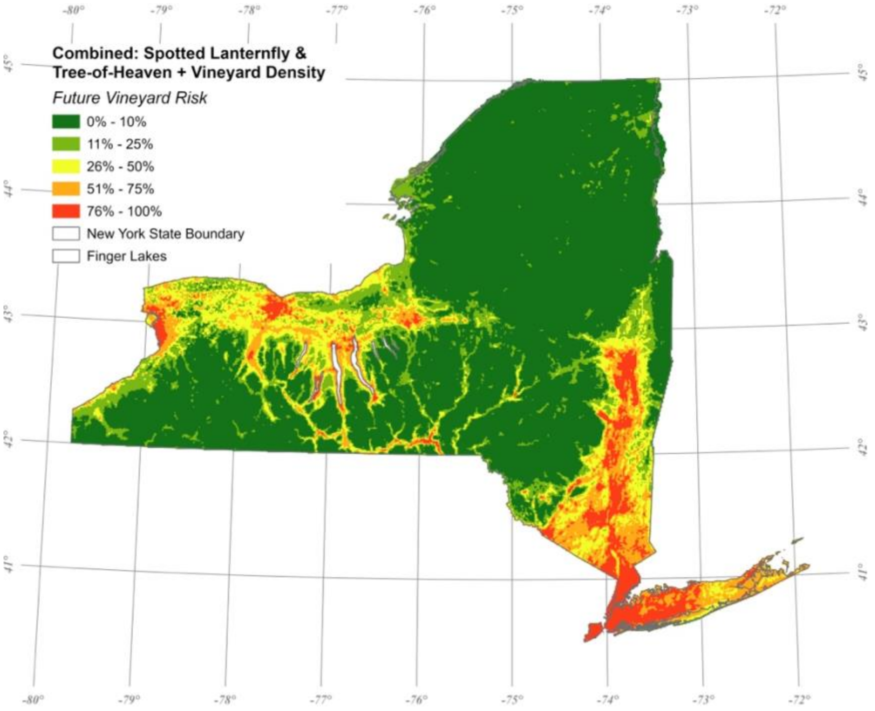

**Figure S9:** A map illustrating vineyard risk under the near-future (2041-2070; RCP 8.5) climate change scenario in New York state. Pixel-wise risk values were determined by applying a vineyard-weighted risk map to the combined (maximum) habitat suitability map of spotted lanternfly and tree-of-heaven. Values were normalized between 0% and 25%.

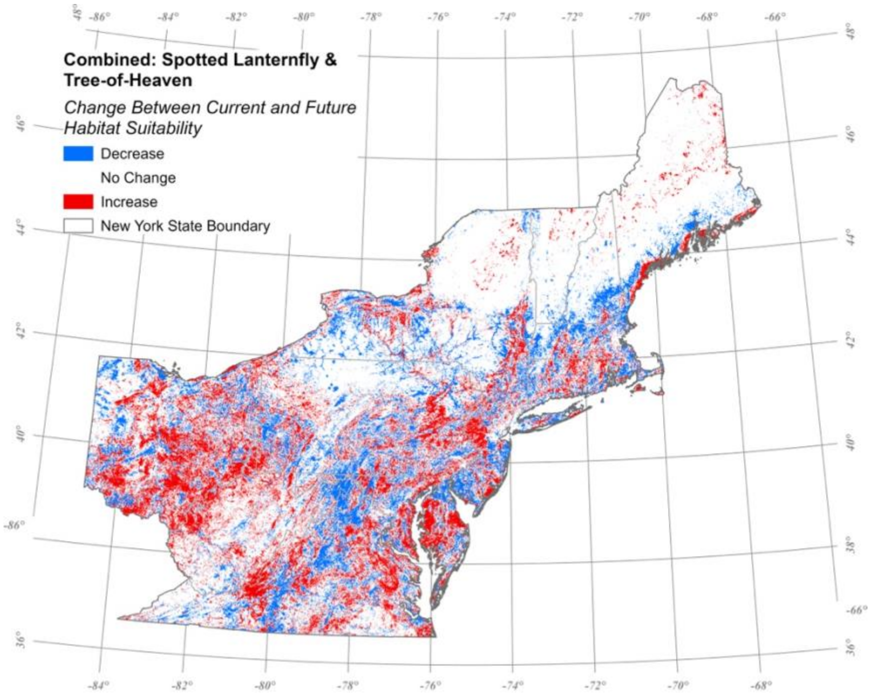

**Figure S10**: A map illustrating the change between the current (2011-2040; RCP 7.0) and near-future (2041-2070; RCP 8.5) combined (maximum) habitat suitability of spotted lanternfly and tree-of-heaven in northeastern United States. States included New York, Pennsylvania, Virginia, West Virginia, Ohio, Maryland, New Jersey, Connecticut, Massachusetts, New Hampshire, Rhode Island, Vermont, and Maine.

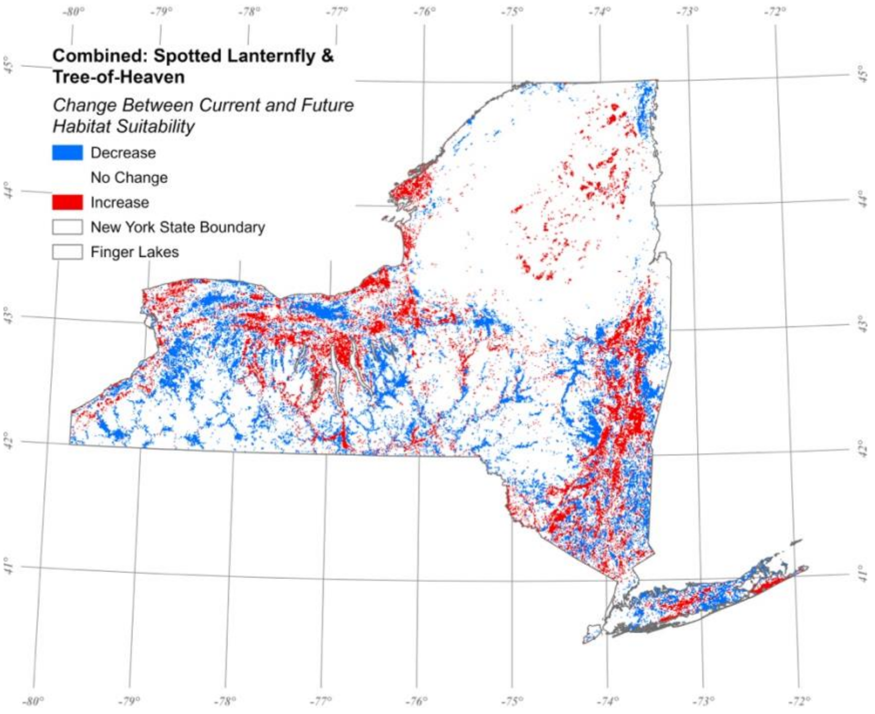

**Figure S11**: A map illustrating the change between the current (2011-2040; RCP 7.0) and near-future (2041-2070; RCP 8.5) combined (maximum) habitat suitability of spotted lanternfly and tree-of-heaven in New York state.
